# Single-cell morphodynamics predict cell fate decisions during mucociliary epithelial differentiation

**DOI:** 10.1101/2025.09.17.676780

**Authors:** Mari Tolonen, Ziwei Xu, Ozgur Beker, Varun Kapoor, Bianca Dumitrascu, Jakub Sedzinski

## Abstract

Cell state transitions underlie the emergence of diverse cell types and are traditionally defined by changes in gene expression. Yet these transitions also involve coordinated shifts in cell morphology and behaviour, which remain poorly characterized in densely packed epithelia. We developed a quantitative live-imaging and computational framework to track thousands of individual cells over time in the rapidly differentiating Xenopus mucociliary epithelium (MCE). From segmentations and trajectories, we extracted dynamic features-cell and nuclear shape, movement, position-to create a time-resolved morphodynamic dataset spanning the full course of differentiation. While single features showed high noise and low separability of ground-truth cell types, supervised machine learning revealed that integrating time-resolved features robustly predicts final cell fate. Gradient-boosted trees and multinomial logistic regression achieved moderate but consistent accuracy, especially for abundant epithelial lineages. Key discriminants included normalized Z-position, membrane-nucleus offset, and absolute experimental time, whereas movement contributed minimally to the results. Our results show that morphodynamic signatures encode predictive information about cell identity and provide a framework linking physical dynamics with molecular state.

## Introduction

Multicellular organisms are composed of a diverse array of specialized cell types, each defined by a distinct phenotype that integrates a specific molecular program with characteristic cell shape and dynamic behaviors such as movement and cell-cell interaction [1,2]. Collectively, these features determine the functional identity of the cell. This integrated profile is often described as the cell state, representing the dynamic manifestation of phenotype and function at a given time [3,4]. Cell states are inherently plastic and context-dependent. Even within homeostatic tissues, individual cells can transition between proliferative and quiescent states, or modulate their behavior in response to mechanical stimuli or intercellular signaling [5–7]. These dynamic cell state transitions highlight the fluidity of cellular phenotypes and their capacity to adapt to physiological demands.

Among the many biological contexts in which cell state transitions occur, development stands out as a particularly striking example, involving large-scale, tightly regulated shifts in cell identity [8,9]. During this process, initially pluripotent cells progressively adopt distinct functional roles through coordinated changes in phenotype. Our understanding of these transitions has so far been primarily focused on molecular aspects, driven by advances in single-cell RNA sequencing, which allows high-resolution profiling of gene expression at the level of individual cells. These efforts have revealed distinct cell states, underlying regulatory networks, and lineage hierarchies that shape developmental trajectories (reviewed by e.g., [10,11]). Furthermore, with the development of computational methods such as pseudotime and trajectory inference, it has become possible to reconstruct the temporal progression of these transitions from static transcriptomic snapshots [12–14].

While recent advances in single-cell ‘omics’ technologies, spanning genetic, epigenetic, and chromatin accessibility profiling, have significantly advanced our understanding of cell state transitions and the emergence of cellular phenotypes, much less is known about the accompanying morphogenetic and behavioral changes that occur during development. Morphology and cell behavior are frequently considered the downstream consequences of fate decisions. However, they may serve as complementary, and potentially predictive, indicators of cell state. For example, the stiffness of the extracellular matrix (ECM) can influence cell fate, demonstrating how mechanical cues, independent of genetic information, can impact cellular morphology and function [15–19]. Unlike molecular measurements, cell morphology is continuously quantifiable in both space and time, making it particularly well-suited to capture the dynamic and spatially embedded nature of cell state transitions, especially when studied through live imaging [20–22].

Image-based single-cell phenotyping has been widely used in biomedical applications such as drug discovery and medical diagnostics, particularly in cell culture systems, where easily quantifiable phenotypes can be efficiently analyzed on a large scale [23–26]. However, its application to developmental biology has been limited, largely due to the technical challenges of processing dynamic, high-resolution imaging data in complex living tissues. Accurately segmenting and tracking cells over time in morphogenetically active environments remains a significant obstacle. Recently, the integration of AI-based methods, including deep learning for segmentation, tracking, and feature extraction, has begun to overcome these barriers [27–30]. Combined with the increasing availability of large-scale, single-cell imaging datasets, including whole-embryo recordings, these advances now make it possible to extend morphodynamic profiling to developmental systems with unprecedented resolution and scale [31,32]. While these studies have begun to relate cellular dynamics to tissue- and organ-level morphogenesis, how cell shape and movement reflect, or even influence, cell state transitions during development remains largely uncharacterized, particularly in densely-packed, multilayered tissues where dynamic morphological features may serve as informative proxies for cell identity and function.

The *Xenopus laevis* mucociliary epithelium (MCE) offers a powerful model for studying cell state transitions within the context of a developing multilayered tissue. This rapidly differentiating, easily accessible epithelium consists of multiple specialized cell types, including multiciliated cells (MCCs), goblet cells, ionocytes (ICs), small secretory cells (SSCs), and basal stem cells. In the fully mature MCE, these cell types are spatially organized into distinct layers: MCCs, goblet cells, ionocytes, and SSCs are located in the superficial (apical) layer, where they carry out mucociliary clearance and secretion, while basal stem cells reside in the basal layer, providing structural support and regenerative capacity [33–39]. In our previous work, we defined the lineage relationships within the MCE and characterized the molecular basis of cell lineage bifurcations, demonstrating that distinct cell fates arise through progressive transcriptional transitions from shared progenitor states [40].

As these molecular programs unfold, cells also engage in coordinated morphogenetic behaviors that shape the developing tissue architecture. These include epithelial thinning, driven in part by epiboly-like movements where cells shift vertically within the epithelium - some moving apically while others are displaced basally [41–44]. A key process in MCE morphogenesis is radial intercalation, through which progenitors of MCCs, ICs, and SSCs insert into the superficial epithelial layer from underlying positions. Notably, the timing of radial intercalation is staggered across cell types, with each population integrating into the surface layer in a temporally distinct manner. The sequential radial intercalation, coupled with apical expansion, ensures proper spatial organization and functional layering in the mature MCE [33,35,41,44]. Together, these dynamic changes in cell shape and movement accompany differentiation, making the *Xenopus* MCE an ideal system to investigate how cell state transitions are defined not only by gene expression, but also by the cell morphology and motility during tissue morphogenesis.

In this study, we developed a quantitative live imaging and computational analysis pipeline to track thousands of individual cells over time in the developing *Xenopus* MCE, capturing dynamic changes in cell shape, nuclear shape, cell position, and movement as cells transition from a multipotent progenitor state to their terminal identities. To explore the connection of single-cell measurements to cell fate, we developed a method to label the five cell types (basal, goblet, ICs, MCCs, and SSCs) after live imaging, and use trajectory information to backtrack cells in their development before specification. We first extracted morphodynamic features from segmentation and tracking data, generating a rich dataset of time-resolved single-cell behaviors spanning the entire differentiation timeline. Despite the biological relevance of the extracted features, individual cell features showed high variability, resulting in a feature space with low separability between cell types. Unsupervised explorations such as PCA revealed limited separation in feature space beyond slight temporal shifts, with the overall distribution of cells remaining largely uniform and intermixed in feature space.

To test whether subtle but consistent differences could nonetheless support fate prediction, we trained a supervised XGBoost classifier using endpoint immunostaining and backtracking-derived ground-truth labels. The model achieved moderate but stable performance across 20 independent stratified train–test splits (80/20), with the highest accuracy for the predominant epithelial lineages: basal cells, goblet cells, and MCCs. Class imbalance was addressed using a combination of undersampling and synthetic oversampling during training. Feature importance analysis revealed that positional and nuclear shape features, especially Z-position and nucleus–membrane offset, were the most informative, while movement features contributed relatively little to cell fate prediction, likely due to collective migration behavior within the tissue. Temporal trends in classification uncertainty and accuracy further showed that morphodynamic signatures of fate emerge progressively during differentiation and often peak prior to terminal maturation.

Together, our findings show that, while instantaneous morphodynamic features are noisy and weakly separable, supervised models trained on time-integrated cell trajectories can extract predictive signatures of cell identity. This study establishes a framework for using dynamic aspects of cell shape and movement, alongside molecular profiles, as complementary descriptors of cell state transitions in dense, developing tissues.

## Results

### A live imaging pipeline for quantifying cell shape and movement during mucociliary epithelial (MCE) differentiation

As a first step toward quantifying how dynamic changes in cell morphology and behavior reflect cell state transitions, we established a high-resolution live imaging and computational analysis pipeline for the developing multilayered *Xenopus* MCE (Figure 1). We used animal cap explants, a well-established *Xenopus* system derived from the superficial ectoderm of the blastula-stage embryo (Figure 1A). When cultured ex vivo under permissive conditions, this tissue undergoes a well-characterized developmental progression from a multipotent state to terminal differentiation, closely mirroring in vivo epidermal development, recapitulating the key morphogenetic events that structure the tissue during embryogenesis (Figure 1B). It reproducibly gives rise to the MCE, generating the full spectrum of epithelial cell types (Figure 1C).

**Figure 1.**
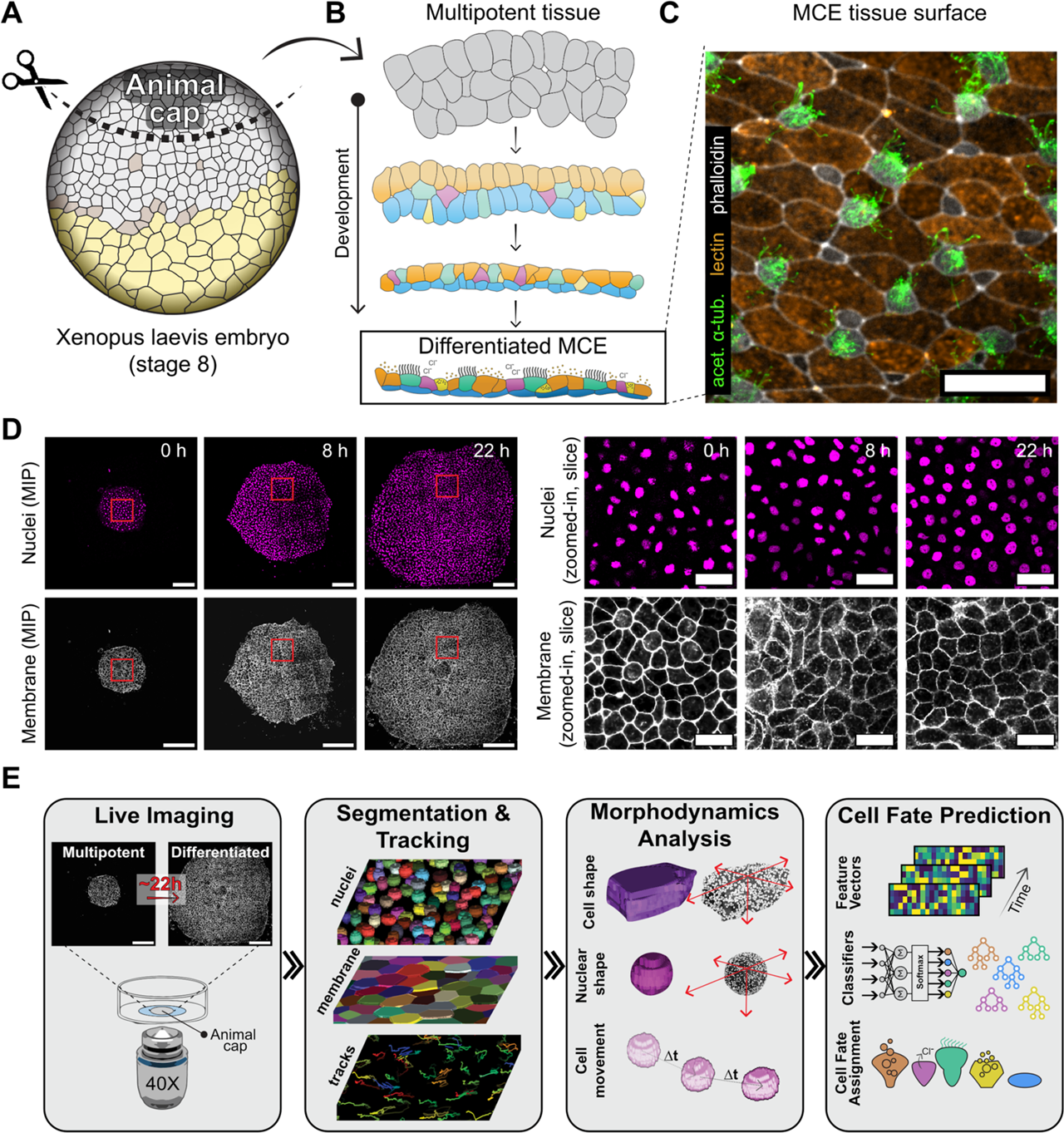
Experimental overview of cell fate prediction. **A.** Animal caps are cut from Xenopus embryos at NF stage 8-9. **B.** The prospective MCE undergoes morphogenetic shaping and cell fate decisions during development into differentiated tissue. **C.** Immunostained differentiated tissue at NF stage 28. acetylated ꭤ-tubulin labels multiciliated cells (green) and lectin labels goblet cells, and small secretory cells in later stages. Ionocytes are distinct for their lack of labeling. **D.** Nuclei and membrane labeling (H2B-RFP and mem-mNeonGreen, respectively) of developing tissue at 0, 8, and 22 h. Overall image scale bar: 300 µm, zoomed-in scale bar: 50 µm. **E.** Image analysis pipeline from live image acquisition to cell fate prediction. Membrane and nucleus objects are achieved via 3D segmentation, and cell shapes and movement are quantified. Single-cell features are then used to classify cell fate by using ground truth data based on cell immunolabeling.

To visualize individual cells over time, we injected mRNAs encoding membrane- and nucleus-localized fluorescent markers (mem-mNeonGreen and H2B-GFP, respectively) into early-stage embryos. Animal caps were dissected at Nieuwkoop-Faber (NF) stage 8-9, while the tissue was still multipotent, and mounted onto fibronectin-coated glass-bottom dishes. Upon adhesion, the tissue flattened, forming a stratified epithelium that enabled continuous optical access throughout development. Volumetric time-lapse imaging was performed over 16–22 hours using confocal microscopy, with z-stacks acquired every 3–5 minutes. This setup allowed us to capture dynamic cell behaviors across the entire course of differentiation, from multipotent states to terminal identities (NF stages 9–30) (Figure 1D, Movie S1). Across three independent experiments, we acquired high-resolution datasets spanning all major developmental timepoints (Table 1).

**Table 1.** Identifying features of cell types for manual annotation based on cell imunolabelling.

| Staining feature | Cell type label |
| --- | --- |
| cell has prominent acetylated α-tubulin staining | MCC |
| cell's nucleus has prominent Tp63 staining | basal |
| cell has prominent and uniform apical lectin staining | goblet |
| cell is small and has granular lectin staining | SSC |
| cell clearly has none of the above | IC |

For analysis, we implemented a custom image processing pipeline to extract morphodynamic features at the single-cell level (Fig. 1E). Segmented cells were tracked across timepoints, and their features – such as shape, size, and movement – were quantified. Using ground truth annotations derived from endpoint immunostaining, we trained classifiers to infer cell fate from morphodynamic profiles (Fig. 1E). Together, this pipeline enables high-resolution, long-term tracking of individual cells as they transition from multipotent to terminal fates within a differentiating tissue context. By capturing both morphological and behavioral dynamics over time, we can begin to map how distinct cell identities emerge during MCE formation.

### Segmentation and tracking pipeline optimized for developing MCE

To characterize single-cell morphodynamics during MCE differentiation, we utilized 3D segmentation and tracking of our volumetric imaging data (Figure 2A). Raw nuclear signals were segmented using a 3D StarDist [45] model trained on manually annotated volumes. Membrane signals were first denoised using CARE [46] and then segmented in 2D on individual z-slices using Cellpose [47,48] (Figure 2B-C, Movie S2). We constructed full 3D membrane objects by assigning slicewise segments across adjacent slices based on maximal pairwise Intersection over Union (IoU), thereby stitching 2D contours into consistent 3D volumes (Figure 2B). This approach reduced spurious objects and improved object consistency along the z-axis. Compared to ground 3D truth annotations, we achieved high segmentation accuracy for nuclei (Table S2). For membrane segmentation, Cellpose produced accurate 2D segmentations with good per-slice performance with a mean per-slice binary Jaccard index of 0.55 ± 0.24 (binary F1 score = 0.68 ± 0.22, Sparse Jaccard index 0.62 ± 0.16, OaGTC Jaccard index 0.72 ± 0.11). However, stitching these slices into coherent 3D cell objects proved more challenging due to variability in cell morphology and limited Z resolution. To evaluate 3D reconstruction quality, we computed intersection-over-union (IoU) using the same predicted and ground truth cell volumes as for the 2D objects. The mean 3D binary Jaccard was 0.55 ± 0.17, (binary F1 score = 0.70 ± 0.15, Sparse Jaccard index 0.48 ± 0.16, OaGTC Jaccard index 0.55 ± 0.10), with lower performance in regions with dense packing or ambiguous membrane signal (Table S2). Despite these limitations, segmentation accuracy was sufficient to support reliable quantification of cell shape and dynamics for downstream analyses.

**Figure 2.**
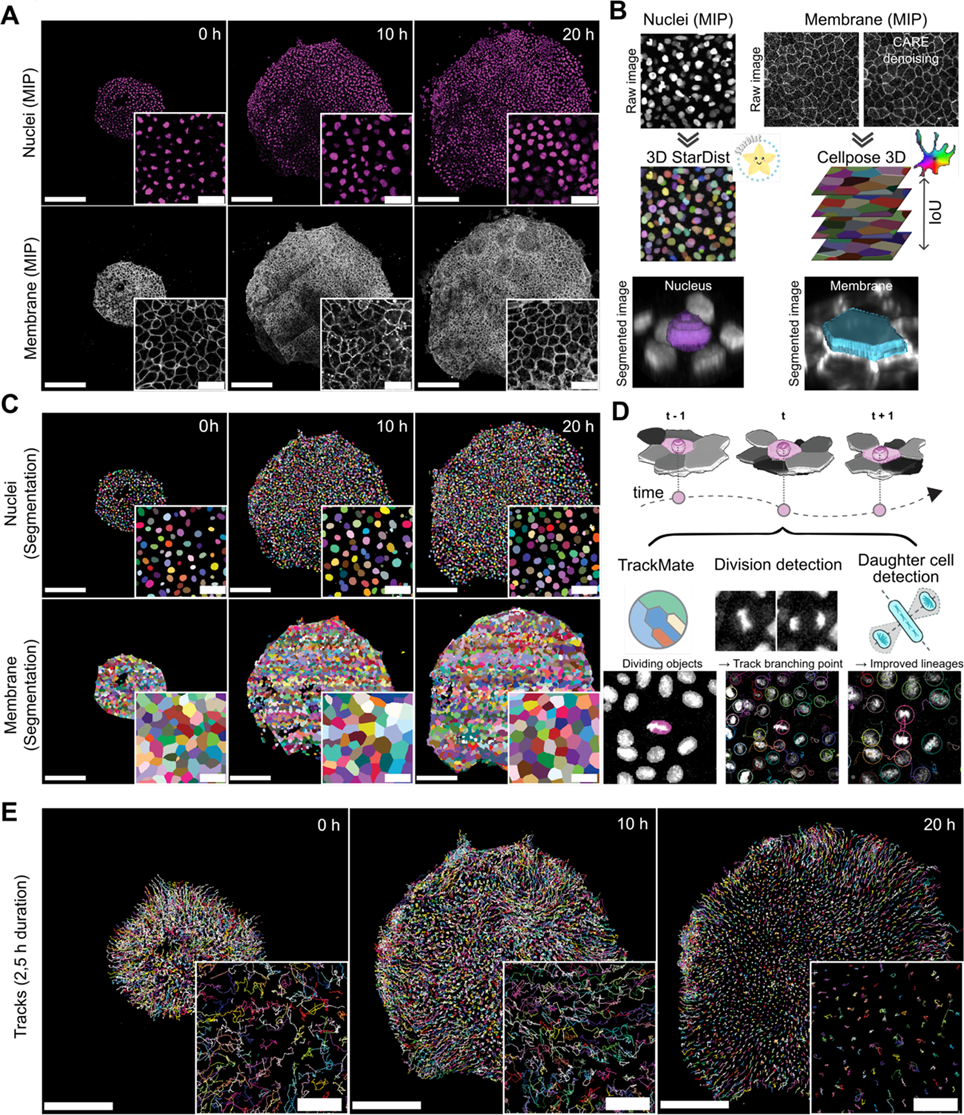
Segmentation and tracking overview. **A-B.** Nuclei and membrane stacks (raw for nuclei, CARE denoised for membrane) are used as inputs for StarDist (nuclei) and Cellpose (membrane). For Cellpose, segmentation is performed slicewise, and 3D objects are reconstructed based on maximal slice-to-slice overlap (IoU). Close-up panels are single z slices (A). **C.** Segmented nuclei and membrane object labels. **D.** Cell trajectories are generated by tracking nuclei objects, which are then assigned membrane objects based on closest object centroid matching. Tracks are initially generated with TrackMate, using StarDist labelled objects as input, and trajectory branching points are imposed at cells that are classified as dividing by Oneat, using a Fiji plugin TrackMate-Oneat. **E.** Resulting cell trajectories at 0, 10 and 20 h. Scale bars for A,C,E: overall image 300 µm, close-up 50 µm.

To reconstruct cell trajectories across developmental time, we tracked segmented nuclear objects, which enabled robust identification of individual cells, even in cases where membrane segmentation was incomplete or missing. To integrate nuclear tracking with shape information, each membrane object was matched to the nearest nuclear centroid in 3D space, thereby linking trajectory data to membrane segmentation (Movie S3). For 3D cell tracking, we used TrackMate 7 [49], which supports labeled nuclear objects as input for spot detection and linking. This approach yielded high linking accuracy (0.919); however, accurate detection of branching events – i.e., cell divisions – remained challenging due to dense tissue packing and limited Z resolution (Table S3). To address this, we integrated Oneat, a custom-built convolutional neural network (CNN)-based division detection module, into the TrackMate pipeline (Figure 2D, Movie S4).

Oneat identifies cell division events as spatiotemporal (TXYZ) coordinates. These coordinates were then used to impose trajectory splitting within TrackMate tracks through our custom-developed TrackMate-Oneat plugin (Figure 2D). The plugin optionally applies the MARI (Mitosis Angular Region of Interest) principle, which filters division events to retain only those in which daughter cells emerge perpendicular to the mother cell’s nuclear major axis (see Materials and methods).

Oneat integration significantly improved mitosis detection compared to TrackMate’s native linking algorithm (TrackMate native branching accuracy = 0.122, TrackMate-Oneat branching correctness = 0.328). By combining Oneat-predicted division locations with trajectory continuity, the system generated more biologically realistic branching structures and reduced the number of false positives typically produced by Oneat alone (Figure S1C–J). However, while applying the MARI principle nearly eliminated false positives, it also led to a reduction in true positive detections (Figure S1E). As such, the user should choose the division detection strategy – Oneat alone or Oneat with MARI filtering – based on the specific goals and tolerance for false positives in their downstream analyses.

Although reconstructing complete cell lineages remains inherently challenging, since even a single linking error can break lineage continuity (Figure S1K), the overall tracking performance was sufficient to achieve our primary objective: generating continuous single-cell trajectories that enable quantification of instantaneous phenotypic features and their integration into complete lineage histories (Movie S5). On average for all datasets, trajectories spanned ≥ 9.5 hours for 33% of cells (of a mean dataset duration of 19.4 h), with the average track length being 7,47 hours, providing robust coverage for downstream analyses of morphodynamics.

### Emerging cell types occupy a continuous and overlapping morphodynamic space

Cell state transitions during development are accompanied by continuous, dynamic changes in morphology and behavior. To quantitatively capture these changes, we defined a *morphodynamic state* for each cell by computing a set of time-resolved morphological and behavioral features, building on methodologies used in recent studies [50–52]. This approach is conceptually analogous to defining a transcriptomic state, where multiple gene expression values are assembled into a high-dimensional vector representing a cell’s identity at a given moment.

For every cell in the dataset, we computed a comprehensive set of features, including spatial position, nuclear and membrane shape descriptors, and instantaneous movement (Figure 3A; Table S4). Segmentation masks were converted into 3D mesh representations and resampled into 1024-point clouds per object (Figure 3C). From these point clouds, we extracted eigenvector-based shape metrics such as surface area, eccentricity and orientation. In addition, for the membrane objects, widely used 2D descriptors of shape were obtained from each 3D membrane object’s center slice (Figure 3B–C; Methods). These measurements were calculated at each time point, forming a high-dimensional feature vector that describes the morphodynamic identity of each cell over time. All feature measurements were calculated at each time point along a cell’s trajectory, forming a high-dimensional morphodynamic feature space.

**Figure 3.**
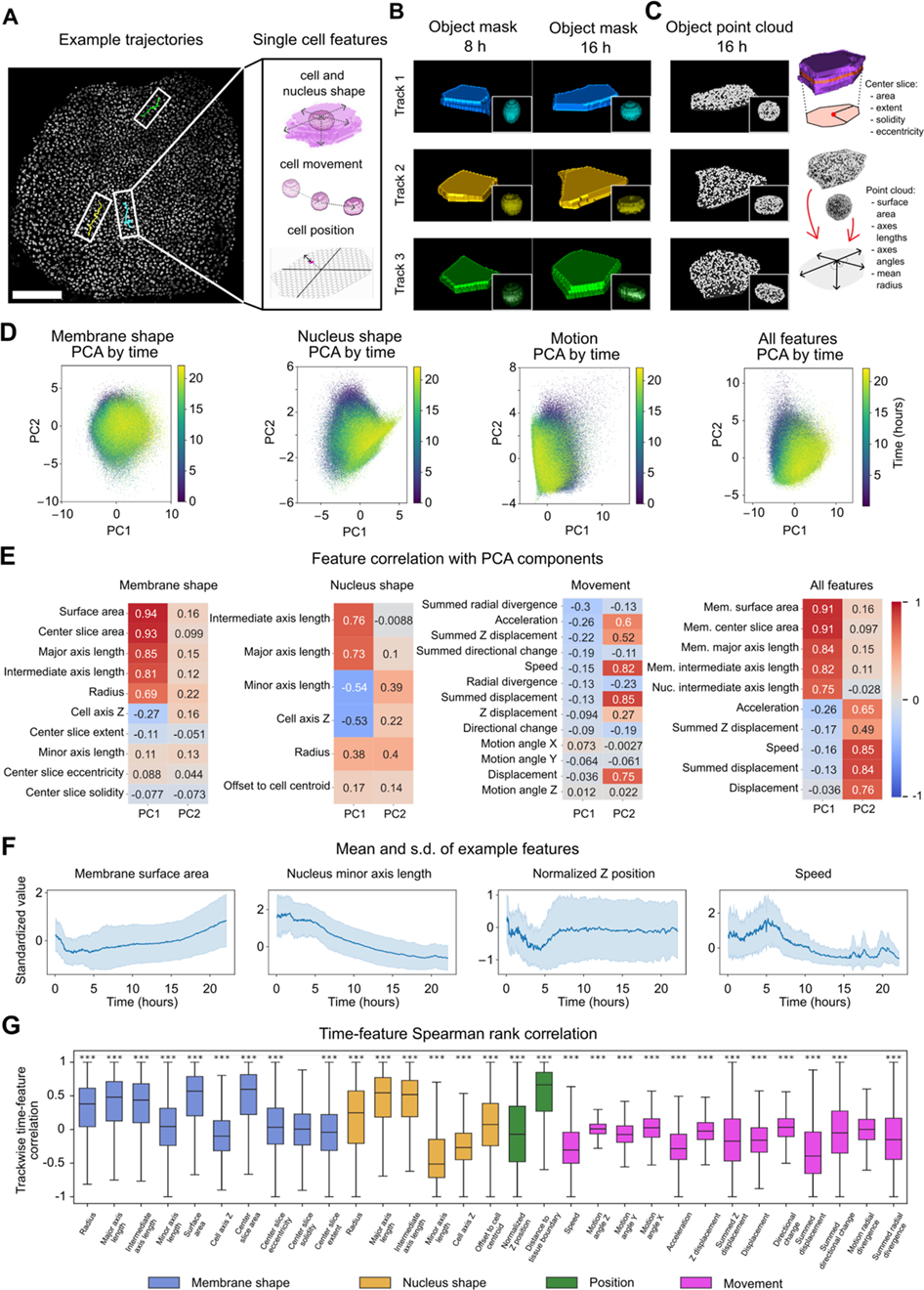
Principal component analysis of single cell features. **A.** Each constructed trajectory in time consists of single cells with shape, nucleus shape, movement, and positional features. Scale bar: 300 µm. **B-C.** Membrane and nucleus shapes of example trajectories at 8 and 16 h, with corresponding point cloud to 16 h. 3D shape measurements of both cell membranes and nuclei are constructed from a point cloud. For membranes, additional 2D shape descriptor features are calculated from the object center slice. **D.** 1st and 2nd principal components of membrane shape, nucleus shape, movement, and all features’ feature space. **E.** Feature correlations with principal components. For all features, only the top 5 correlating features for PC1 and PC2 are shown. **F.** Selected feature trends across developmental time for all cells. **G.** Spearman rank correlations of each feature with time, calculated per trajectory, and a boxplot shows the distribution of correlations. Asterisks denote features with significant nonzero correlation after FDR correction (*** = *p* < 0.001), (Wilcoxon signed-rank test across trackwise correlations).

Principal component analysis (PCA) was used to reduce dimensionality across nuclear shape, membrane shape, and motility features, both individually and in combination with positional features (Figure 3D, Figure S2C). PCA was performed on a standardized feature matrix comprising n = 4269 cell trajectories and d = 34 features (Table S4). For feature space fit with all features, the first three principal components captured 37.3% of the total variance (PC1: 17.6%, PC2: 10.7%, PC3: 8,9%). However, PCA projections did not reveal distinct clusters corresponding to specific cell types or developmental stages. Instead, cells were distributed continuously in feature space, with slight shifts over developmental time. These results suggest that morphodynamic transitions occur along a spectrum, rather than through discrete jumps between feature space groupings. These observations were consistent across datasets (Figure 3D, Figure S2A, C), reinforcing the notion that cell state transitions in the MCE are gradual and multidimensional, without sharp boundaries in morphodynamic space.

Correlations between individual features and principal components were used to interpret the major axes of variation (Figure 3E), while time-resolved plots of selected features highlighted global trends in standardized features, such as decreasing motility and progressive cell flattening over time (Figure 3F). However, most individual features exhibited limited temporal variation and failed to show significant trends over time (Figure S3). When individual cell trajectories were visualized in feature space, there was no consistent temporal progression, except for positional features, which showed gradual changes over time (Figure S2B–C). This was further supported by Spearman rank correlations between feature values and time, which revealed statistically significant, albeit low, correlations for the majority of features (Figure 3G). Thus, unlike shape-based profiling in cultured cells, where morphologies are highly heterogeneous [22], cells within this densely packed 3D epithelium exhibit more constrained morphologies and movement behaviors. These similarities mean that, when all cells are placed in a shared feature space, they are not easily distinguishable based on shape or movement features alone.

### Establishing a ground-truth dataset of cell lineage histories with backtracking

Given the limited ability of unsupervised approaches to predict cell fate from morphodynamic features alone, we shifted toward a supervised strategy that incorporates lineage history. To enable this, we first established a ground truth dataset by assigning final cell identities through endpoint immunostaining and morphology-based annotation of individual lineages. Following completion of live imaging, samples were fixed and immunostained for four well-characterized epithelial cell type markers corresponding to MCCs, goblet cells, small SSCs, and basal cells, as previously described. Cells lacking marker expression but exhibiting characteristic radially intercalating cell morphology and position were manually annotated as ICs (Figure 4A-B). This endpoint labeling enabled us to assign final fates to approximately 75% of all automatically tracked cell trajectories.

**Figure 4.**
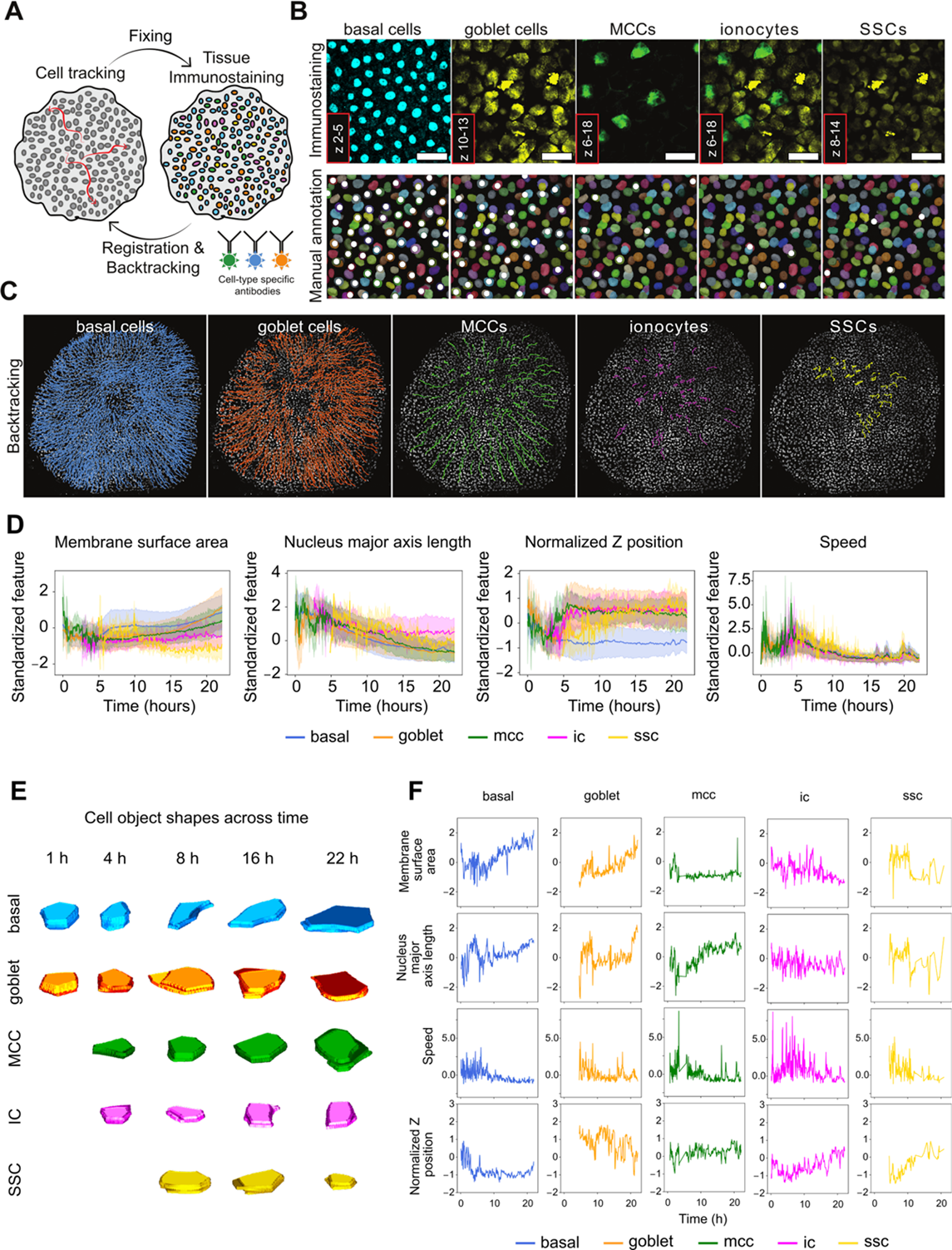
Backtracked cell types’ feature distributions. **A-B.** Live imaged cells are fixed at the end of the experiment and immunolabelled for MCE cell types (A). The cell fate is assigned to a track based on manual annotation of the final timepoint cell type based on the presence of immunolabel in a cell (B). **C.** Tracks with cell fate labelling, overlaid against the last timepoint. **D.** Mean and s.d. of selected features, colored by cell type. **E.** Membrane shape of randomly selected cells, 1 per cell type, at 1 h, 4 h, 8, 16 h and 22 h. **F.** Single cell features of randomly selected cells across time, 1 per cell type.

To ensure representative sampling, we manually curated trajectories corresponding to MCCs (n = 156), ICs (n = 45), and SSCs (n = 11), which represent less abundant and morphologically more variable cell types. In contrast, trajectories for basal (n = 1457) and goblet cells (n = 909), which are more abundant and morphologically homogeneous, were initially annotated automatically, with manual correction of randomly selected subsets (n = 54 and n = 10 for basal and goblet cells, respectively) used for subsequent classifier training (Figure 4C, Movie S6). This annotation resulted in a high-confidence ground truth dataset comprising 2576 total trajectories for subsequent classification tasks (see below).

To evaluate whether individual features were potentially predictive of final fate, we first examined the temporal dynamics of selected morphodynamic features across cell types (Figure 4D). Consistent with earlier observations from the global feature space, single features alone provided limited discriminatory power between cell types, showing high within-type variability and overlapping distributions (see also Figure S5).

To more comprehensively assess whether combinations of features could distinguish cell types, we projected our ground-truth backtracked trajectories into principal component spaces derived from nuclear shape, membrane shape, motility, and positional features, as in Figure 3D. As before, cells did not form clearly separable clusters by fate in PCA space (Figure S4B). This observation held even when trajectories were divided into developmental time windows and analyzed separately by cell type (Figure S4C), indicating that feature divergence was subtle and gradual over time.

To quantify the seemingly low separability, we computed unsupervised clustering metrics including Adjusted Rand Index (ARI), Normalized Mutual Information (NMI), and Average Silhouette Width (ASW) across multiple feature combinations. Scores were uniformly low (mean ARI = 0.019 ± 0.001, NMI = 0.053 ± 0.001, ASW = -0.156 ± 0.004), with positional features contributing most to weak separability (mean ARI = 0.104 ± 0.004, NMI = 0.133 ± 0.003, ASW = -0.118 ± 0.005), and scores improving over developmental time (Figure S4A). Similarly, we trained a K-nearest neighbor classifier (KNN, k = 5) using morphodynamic features to predict final cell fate. The mean accuracy of the KNN classifier trained on all features remained at 65.8% across 5 folds (baseline = 54.0% for stratified dummy classifier), with balanced accuracy score at only 28.1% against a baseline of 24.8%, confirming the limited predictive power of unsupervised or shallow models in the absence of lineage context (Figure S4A).

Despite this, visualizing selected individual cell trajectories revealed temporally dynamic shape and movement changes, with subtle but fate-associated trends (Figure 4E–F, Movie S7). While shape and movement trends show dynamic changes across time, normalized Z position proves to carry the clearest single-cell signature of cell type, with basal cells staying stably low after early variation, goblet cells having a descending trend, and ICs, MCCs, and SSCs cells having an ascending trend (Figure 4F). These observations underscore the need for supervised, temporally aware models to robustly link early morphodynamic behaviors to eventual fate.

### Morphodynamics-based multiclass classification of cell fate using XGBoost

To assess whether combinations of morphodynamic features could jointly predict cell fate, we trained multivariate multiclass classifiers using XGBoost and a multinomial logistic regression model from scikit-learn (Figure 5A) [53,54]. These models use ensemble learning (XGBoost) or linear decision boundaries (logistic regression) to predict final cell fates based on morphodynamic feature vectors. Each cell at each time point was treated as an independent observation, and models were trained to predict endpoint fates derived from immunostaining-based backtracking. All cells in each track were thus assigned a cell type based on their final cell fate, as we observed through manual track annotation that daughter cells in the same trajectory end up with the same final cell fate (data not shown).

**Figure 5.**
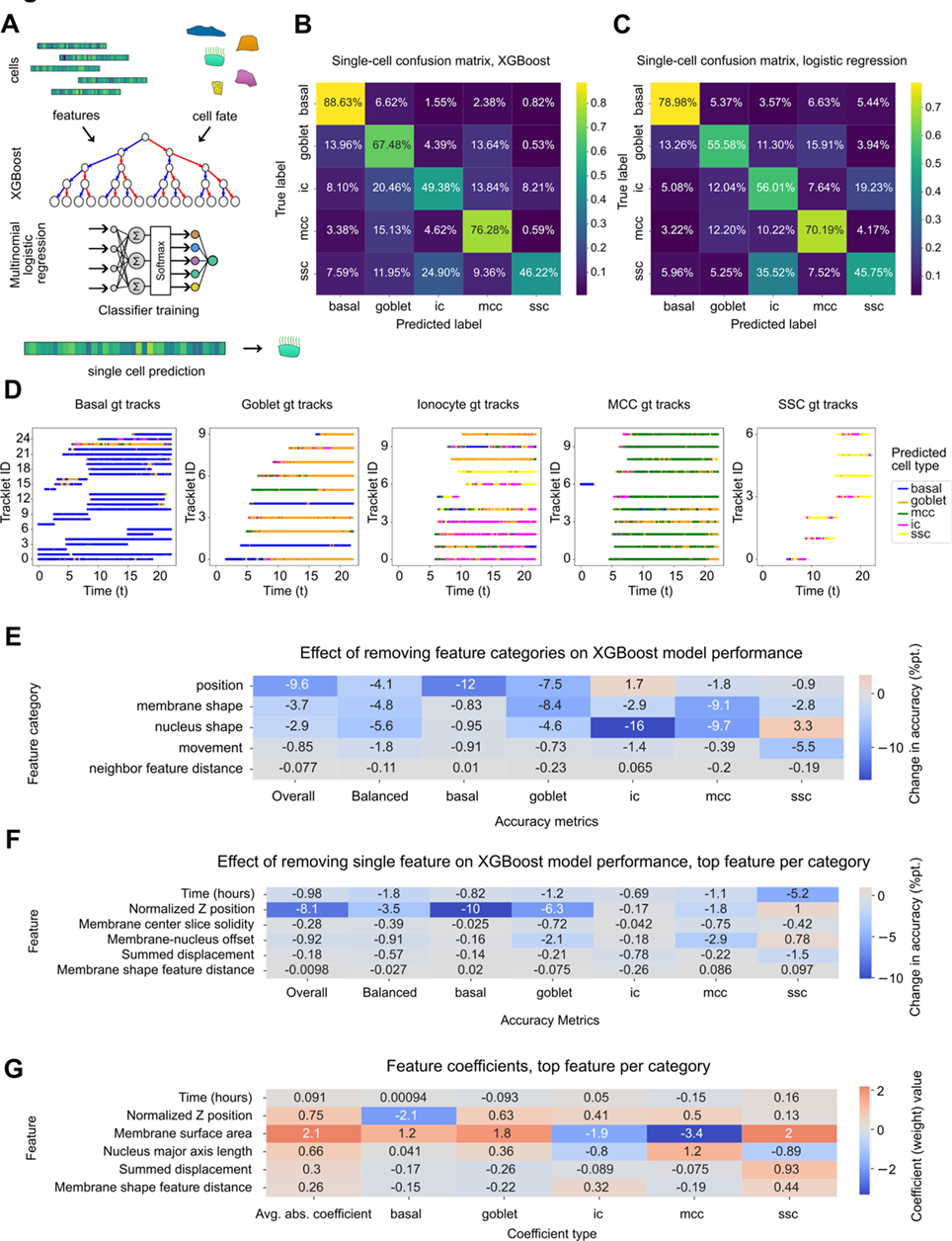
Classifying cell fate using XGBoost and logistic regression. **A.** Feature vectors of single cells are used as independent features, and cell fate as a dependent feature, for training a classifier model, based on either XGBoost or multinomial (softmax) logistic regressor. Then, each cell in the test dataset is assigned a cell fate based on its feature vector. **B-C.** Confusion matrices of test dataset cell fate prediction vs. ground truth, for XGBoost model and logistic regression model, respectively. **D.** Example test dataset trajectories showing single-cell predictions per track. **E-F.** Percent point-wise accuracy difference to baseline accuracy of the XGBoost model for leaving out a feature category (E) or a single feature (F, only the top feature per feature category is shown). **G.** Feature coefficients for a logistic regressor model, only the top feature per category is shown. All calculations in B, C, E, F, and G are averaged out for 20 iterations of randomized test-train split, model training, and prediction.

In addition to morphodynamic features, we included absolute experimental time as an input variable, which improved prediction performance by 0.98 percentage points on total accuracy and 1.8% percentage points on balanced accuracy (Figure 5F) of the XGBoost model. Model performance was assessed over 20 independent random 80/20 train–test splits, stratified by cell type and time. Across these iterations, both models achieved similar performance: XGBoost reached a mean accuracy of 80.6% (±0.2%), balanced accuracy of 61.4% (±1.2%) and macro F1-score of 0.56, while logistic regression achieved 70.3% (±0.3%) accuracy, 62.9 (±1.2%) and a macro F1-score of 0.45 (Figure 5B–C). Prediction performance was highest for basal (F1 (XGBoost) = 0.90, F1 (logistic regression) = 0.85), goblet (F1 (XGBoost) = 0.72, F1 (logistic regression) = 0.64), and MCC (F1 (XGBoost) = 0.56, F1 (logistic regression) = 0.45) lineages, which are the major epithelial sublineages in the MCE. Minority classes, including ICs (F1 (XGBoost) = 0.34, F1 (logistic regression) = 0.27) and SSCs (F1 (XGBoost) = 0.25, F1 (logistic regression) = 0.06), remained difficult to classify, despite application of class balancing via a combination of SMOTE-based synthetic oversampling and undersampling (Figure S6A).

To evaluate how well classifier predictions reflected the overall differentiation process, we visualized single-cell predictions across trajectories (Figure 5D). For basal, goblet, and MCC lineages, most predictions along a given trajectory consistently matched the true fate label, indicating that morphodynamic signals are relatively stable and informative at the trajectory level. These data suggest the potential for future models that incorporate full trajectory information for improved classification accuracy.

We next examined how classifier confidence and accuracy evolved over time. As expected, both metrics increased during development for most cell types, consistent with progressive fate specification and increased classwise morphodynamic separation (Figure S6B–C). For example, the mean classifier confidence across all cell types rose from 0.80 at ∼5 hours to 0.89 by ∼22 hours, accompanied by an increase in single-cell predictions’ class-balanced accuracy from 50.4% to 75.1%.

Interestingly, prediction confidence and accuracy for MCCs peaked at ∼15 hours (mean confidence = 0.82; accuracy = 80.6%) and declined at later stages (to 0.78 and 57.1%, respectively, by 22 hours). These results suggest that MCCs reach a morphodynamic plateau before the final timepoints, after which increasing overlap with goblet cell features may lead to ambiguity or phenotypic convergence (Figure S6C).

Early in development, predictions were heavily biased toward the basal cell class, which is both the most abundant and most consistently tracked from the start of imaging. This bias likely arises from both biological and technical factors: basal cells reside in the lower epithelial layer, which is most visible early in development, while other lineages become detectable only after tissue flattening improves imaging of apical cells.

To understand which features contributed most to prediction, we assessed feature importance via two complementary methods: leave-one-feature-out accuracy drops for XGBoost (Figure 5E–F, Figure S6D) and coefficient magnitudes for the logistic regression model (Figure 5G, Figure S6E). Across both models, positional features were consistently among the most informative. In particular, normalized Z-position had the greatest predictive value, likely reflecting known apico-basal differences between cell types. Removing Z-position caused the largest performance drop in both models.

Interestingly, time was also a strong predictor, especially for SSCs, which emerge later in development. Excluding time as a feature led to a notable decrease in SSC classification accuracy, while only minimally affecting predictions for more abundant lineages (Figure 5F, Figure S6D).

Among morphological features, nucleus–membrane centroid offset emerged as the most predictive individual variable in the XGBoost model. In contrast, membrane surface area and major axis length were most influential for the logistic regression model (Figure 5F, G). These differences likely reflect how the two models capture feature interactions: XGBoost considers non-linear combinations, while logistic regression relies on additive linear effects.

Overall, nuclear shape features were comparatively informative to membrane shape features, even though to the human eye, cell types are more distinguishable by their membrane shape. This result is likely due to higher segmentation fidelity of nuclei. Conversely, movement features – both instantaneous and time-aggregated – contributed the least to model performance (Figure 5F). These results align with earlier observations that movement patterns in the developing epithelium are largely collective and not fate-specific.

Together, these analyses demonstrate that multivariate morphodynamic features contain predictive information about cell fate, especially when incorporating positional context and developmental time. However, classification performance remains limited for minority populations, and trajectory-level or temporally contextual models may be needed to fully resolve early fate specification.

## Discussion

In this study, we present a full experimental and computational pipeline for imaging, segmenting, tracking, and classifying single-cell morphodynamic behaviors during *Xenopus* mucociliary epithelium (MCE) development. We capture nucleus and membrane dynamics of single cells over time and link them to terminal cell fates via immunostaining, thus generating a continuous, high-dimensional phenotypic characterisation of *Xenopus* embryonic MCE development. Our work provides a foundation for understanding how cell fate is linked to dynamic changes in shape, position, and movement within a complex, densely packed, and actively developing tissue.

### Cell morphodynamics as a continuous phenotype

Our study supports the idea that cell fate can be partially inferred from morphodynamic features alone. However, unlike in cultured cell systems – where shape-based classification of cell states can impressively resolve distinct drug treatments and cell types [26], our in vivo data shows single cells exist in a highly overlapping but noisy feature space, especially early in development. These results are consistent with observations in other complex tissues, where unsupervised clustering based on cell shape features shows initial promise but continues to face significant challenges in reliably distinguishing cell types; In the anterior visceral endoderm, cell fate is still largely position-dependent despite regional shape (and behaviour) trends [55], and in the inner ear, nucleus and cell shape could separate hair cells, but struggled to resolve supporting cell subtypes [56]. Together, these studies underline how densely packed, morphogenetically active tissues pose challenges for inferring fate from morphology alone, partly due to the limited resolution of 3D tissue imaging, partly due to cell shape similarity in packed tissues.

Despite these challenges, our classifier models are able to extract meaningful signals of phenotypic state, especially when observing the time-resolved predictions across trajectories. Our data mirror developments in single-cell transcriptomics, where pseudotime helps supplement the understanding of the developmental timeline, when the data is based on stagewise snapshots [40]. Trajectory-resolved phenotypic data could complement transcriptomics approaches, as it provides a more continuous view of cellular state transitions over time [57]. Unlike scRNA-seq, which captures static snapshots without spatial context, or spatial transcriptomics, which may achieve cellular resolution, but at the expense of transcript depth, morphodynamic tracking provides dense temporal sampling with rich positional information embedded in tissue context. As such, it has the potential to fill in the temporal and spatial gaps in transcriptomics-based fate mapping, offering a new, highly data-rich aspect to the “omics” view of tissue differentiation.

### Comparisons to related approaches

The efforts to track dynamic, developing 3D tissues in a high-throughput way are continuously being developed [28,31,58,59]. On top of that, linking these tracks to single-cell fates to provide large-scale lineage tracing of tissues is still challenging. While lineage tracing embryos and embryonic tissues has been used to map single-cell fates (e.g., [60,61]), these efforts have not focused on detailed shape/movement data. Our approach emphasizes quantifying cell morphogenesis over time, offering new insights into how the combination of position, shape, and movement may predict fate decisions.

Similarly, while tracking of cell phenotypes via cell shape and movement has been recently studied in vitro [25,52,62], our pipeline demonstrates that even in complex tissues, morphodynamic phenomics can be scaled and analyzed like other ’omics’ approaches, thus proving the potential of the addition of “phenomics” data, recently demonstrated by [55].

### Limitations and data constraints

Our segmentation and tracking pipeline is optimized for large-scale throughput, which can lead to incorrect lineage reconstruction or assignment, especially during divisions or in less optically accessible regions. This likely biases our dataset toward better-segmented and longer-lived trajectories, such as basal cells. While our segmentation and tracking pipeline is optimized for large-scale throughput, challenges remain in accurately reconstructing lineages, particularly during cell divisions or in regions with limited optical accessibility. These limitations may introduce bias toward trajectories that are more easily segmented and tracked—such as those of basal cells. To mitigate this, we invested substantial effort into refining segmentation algorithms, generating manually annotated ground truth data, and performing targeted manual corrections. As a result, our dataset offers a high-quality, representative view of diverse epithelial cell types. These efforts also establish a robust framework for generating reliable, high-throughput morphodynamic data in future studies with improved scalability and accuracy.

Our data underscore a central challenge in modeling morphodynamic phenotypes: cell types often fail to form distinct clusters in feature space, particularly at early developmental stages, making unsupervised methods insufficient for fate identification. Consequently, while instantaneous morphology may carry some information about cell state, fate becomes apparent only gradually through changes in shape, position, and movement. As a result, effective classification requires supervised models capable of learning more confined, time-dependent patterns – like CNNs or XGBoost – rather than relying on distinct feature space separation as in omics-based analyses. Our model achieves reasonable fate prediction despite the overlap in feature space, suggesting that morphodynamic features carry consistent temporal signals, even if they are not easily visualized through unsupervised clustering efforts.

### Broader implications and future directions

This work highlights the potential of a morphodynamic single-cell phenomics framework for studying live, developing tissues. By demonstrating that time-resolved cellular features can predict cell fate with meaningful accuracy, it underscores the power of combining high-throughput imaging with advanced computational analysis to investigate developmental mechanisms, tissue self-organization, and cellular responses to perturbation.

Importantly, our quantitative pipeline can be readily extended to other systems or combined with molecular perturbations. For instance, integrating morphodynamic data with single-cell transcriptomics or targeted perturbations of key signalling pathways could reveal causal links between gene expression, developmental signalling, and cell phenotype. Moreover, because the model generates time-resolved predictions for each cell based primarily on shape and position, it could also enhance tracking accuracy by optimizing for continuous cell fate prediction.

While single-cell morphodynamic features in tightly packed epithelial tissues may lack strong separability on their own, future strategies could exploit the inherent spatial patterning of developing tissues. Integrating information about neighboring cells and local context, such as differences in features relative to adjacent cells, patterns of cell–cell interaction, or local geometry, may reveal discriminative signatures that are obscured when analyzing cells in isolation. Such spatially informed models have the potential to substantially improve predictive power and yield deeper insights into the collective dynamics of tissue development.

Ultimately, this approach moves toward a more dynamic view of development, where cell fate is not just defined by signalling pathways but a continuously changing trajectory through shape, movement, and position in a tissue context.

## Methods

### mRNA synthesis

To transcribe mRNAs for fluorescent protein expression, pCS2 or pCS2+ plasmids were linearized with NotI-HFR (NEB), purified by a gel extraction kit (Qiagen), and then used as templates for in vitro transcription using the mMachine SP6 kit (Ambion). Synthesized mRNA was purified by LiCl precipitation, dried, and dissolved in RNase-free H2O (Ambion)

### *Xenopus laevis* husbandry and embryo manipulation

Wild-type X. laevis were obtained from Nasco, Wisconsin, Fort Atkinson, WI, USA. The Danish National Animal Ethics Committee has reviewed and approved all animal procedures under permit number 2017-15-0201-01237. Ovulation was induced in X. laevis adult females by injecting 500 U/animal of Human Chorionic Gonadotropin (Chorulon). Eggs were harvested and fertilized in vitro using macerated testes from male frogs in ⅓x Marc’s Modified Ringer’s (MMR) solution. After 2 hrs, fertilized eggs were dejellied with 3% cysteine (pH 8.0) solution. Cleaving embryos were then washed and reared in 1/3× MMR solution. For mRNA microinjection, embryos were transferred to a 2% Ficoll in 1/3× MMR solution at the 4-cell stage and injected in both ventral blastomeres twice with 100 pg H2B-RFP and 50 pg 3xGFP-E-Cadherin and 100 pg GFP-C-Cadherin or 100 pg mem-mNeonGreen encoding mRNA constructs, 10 nl per injection. All RNA constructs were diluted in RNase-free H2O (Ambion). H2B-RFP was used in all experiments to label nuclei, and coinjected with either mem-mNeonGreen or a combination of 3x-GFP-E-Cadherin and GFP-C-Cadherin for membrane labelling. After injection, embryos were incubated at 13°C until NF stage 8-9 for animal cap cutting.

### Animal cap dissection

Fibronectin (FN) coverslips were prepared by pipetting a 100 µl droplet of 100 µg/ml bovine plasma fibronectin (Sigma-Aldrich) on a 25 mm 1.5 H coverslip (Superior Marienfeld) and incubating o/n RT. Animal caps were cut from dechorionated embryos in 1/3x MMR, washed in Danilchik’s for Amy medium (DFA), and transferred into an imaging chamber fitted with the FN-coated coverslip in DFA. To allow for tissue attachment to the coverslip, animal caps were placed deep cell layer towards fibronectin, affixed using a coverslip chip and silicone grease, pressing the tissue down lightly for approximately 1 hour. After attachment, the coverslip chip was removed, and live imaging was set up immediately.

### Confocal microscopy and live imaging setup

Images of explanted animal caps were acquired with confocal laser scanning inverted microscopes Zeiss LSM880 and LSM980 with Airyscan2 detector equipped with a ×40 C-Apochromat W autocorr M27 water immersion objective (NA = 1.2, working distance = 0.28 mm) (Carl Zeiss Microscopy). All imaging was performed at room temperature, and images were acquired in the regular confocal mode, using excitation lasers of 488 nm for mem-mNeonGreen or 3x-EGFP-E-Cadherin and GFP-C-Cadherin, and 561 nm for H2B-GFP. The obtained tiles were stitched using ZEN Black software (Carl Zeiss Microscopy).

Timelapse imaging was performed using Experimental Designer mode in ZEN Black software to allow for multiple tile and time interval setups. To accommodate for tissue growing in the field of view during the experiment, images were acquired using 3x3 tiling at experiment time ∼0-5 h, 4x4 tiling at ∼5-10 h, and 5x5 tiling until experiment end at 16-22 hours. Tiles were acquired with a 10% overlap for all tilings. Time interval was chosen according to the tiling to be as fast as possible to enable reliable tracking of cells, using 120 sec, 200 sec, or 300 sec time intervals, respectively. An optical resolution of 0.69 x 0.69 x 2.0 µm/pixel (XYZ) was used for all experiments.

### Post-live imaging immunostaining

After live imaging, DFA medium was aspirated and 2 ml of 3,7% PFA in 1X PBS was added directly to the imaging chamber. Tissue fixing was monitored by taking confocal images with the previously mentioned settings. The sample coverslip was transferred to a plastic petri dish, and fixing was continued for a total of 30 min. Immunostaining was further continued as explained in [63], except for omitting sample H_2_O_2_ bleaching. Primary antibodies used were specific for acetylated-α-tubulin (from rabbit, 1:400, Cell Signalling Technologies), and Tp-63 (from mouse, 1:200, Abcam). Secondary antibodies used were anti-rabbit Alexa-488 or Alexa-647 (from goat or donkey, 1:400, Thermo Scientific) and anti-mouse Alexa-568 or Alexa-647 (from goat, 1:400, Thermo Scientific). Simultaneously with secondary antibodies, samples were incubated in 1:1000 lectin PNA Alexa 568 or Alexa 488 (Thermo Scientific) and 1:1000 DAPI (Thermo Scientific). The sample was placed back in an imaging chamber in 1X PBS and imaged using the same settings as the last timeframe of live imaging, adjusting for 4 channel imaging using a combination of 405 nm, 488 nm, 561 nm, and 639 nm excitation lasers, and setting Z optical sectioning to 1 µm.

### Image processing overview

The image processing pipeline is shown in Figure 1, and details are discussed in paragraphs below. In short, tiled timelapse images were exported as 8-bit TIFF files, combined into a timelapse by matching 3 x 3 and 4 x 4 tiled image canvas size with 5 x 5 tiled image size on ImageJ. Z stacks were tilt corrected using a custom ImageJ macro, and if notable drift was present for the dataset, XYZ drift was corrected using the Fast4DReg ImageJ plugin [64]. Membrane channel images were denoised using a custom-trained CARE model in the CSBDeep Python package [46]. Cell nuclei on the nuclei channel were segmented using a custom-trained 3D StarDist model. 3D cell membrane volumes were constructed by segmentation using a custom CellPose 2D mode on the membrane channel, after which 2D slices were stitched into 3D objects using a custom Python script. Segmented nuclei were tracked using TrackMate LAP linker, and trajectories were further corrected by enforcing trajectory branching points using the TrackMate-Oneat plugin on ImageJ. Trajectory data was further processed using the NapaTrackMater Python package, combining cell lineage history with single cell shape (based on 3D measurements of segmented nucleus and cell membrane objects), and movement and tissue position (based on nuclei objects XYZ coordinates).

### Nucleus segmentation

To segment nuclei in 3D, we started out by training a custom model using the StarDist 3D framework [45], optimized for instance segmentation of star-convex shapes in volumetric data. For training, data for annotating 3D nuclear masks were selected from representative ROIs across different developmental timepoints. Augmentations were applied to training patches prior to training. We integrated StarDist segmentation into our VollSeg package, which first finds tissue region of interest from raw nucleus data based on UNET segmentation of Z maximum projected nucleus channels per timepoint, reducing the data input for prediction, and improving pre-segmentation normalization step on images where tissue area takes up a small portion of the whole image area.

The trained model was applied to all volumetric datasets to segment nuclei in the nuclear channel using the VollSeg framework. To assess segmentation quality, we compared predicted nuclear masks to ground truth manual annotations. All nuclear segmentations were performed in batch using GPU acceleration via TensorFlow.

### Membrane segmentation

We trained a Cellpose neural network model by utilizing their human-in-the-loop training workflow [47,48], using their existing ‘cyto2’ model as a backbone. To generate a training dataset, representative 2D ROIs were selected across different timepoints in developmental time. These frames were manually annotated to generate training data. Both labelling and training were performed on CellPose GUI to generate a custom CellPose model, which was then used on the experimental datasets, using a flow threshold 1.0, a cell probability threshold of -1.0, and a cell diameter of 33.2. We stitched 2D slices into 3D objects using the IoU stitching method from CellPose using IoU threshold of 0.6, post-processing each achieved 3D object by using the initial objects’ center slices as object seeds, assigning the rest of the 2D slices to the seed with the highest IoU, maximising overlap between consecutive slices. Furthermore, residual objects of thickness below 2 slices were discarded. Every image dataset’s membrane channel was batch-segmented using the custom model via the Cellpose command-line interface through PyTorch, after which above mentioned postprocessing was applied.

### Segmentation quality measures

We evaluated the segmentation performance of the nucleus and membrane using the F1-score of binarized images, and three variations of the Jaccard Index. The standard Jaccard Index was computed on binarized masks as the ratio of intersection to union over all foreground pixels. The Sparse Jaccard Index matched individual ground truth and predicted objects by computing pairwise intersection-over-union (IoU) scores only for label pairs with overlapping pixels, then averaging the best non-zero IoUs for each ground truth object. The Object at Ground Truth Centroid (OaGTC) Jaccard Index measured the IoU between each ground truth object and the predicted object located at its centroid, averaging the results across all ground truth objects with valid matches. Calculations were averaged across multiple annotated ground truth images (Table S2).

### Tracking quality measures

Automated tracking performance (without or with cell division correction) was first benchmarked against automatically generated, manually annotated (“silver” ground truth) cell tracks on a data ROI (Figure S1A) using the Python package ‘ctc_metrics’, which implements standardized Cell Tracking Challenge (CTC) metrics.[65,66] We computed key measures such as detection accuracy (DET), tracking accuracy (TRA), association accuracy (LNK), cell divisions (CT), mitotic branch correctness (BC), and additional metrics like false positives/negatives (TF) and cell cycle annotation accuracy (CCA).

Additionally, to evaluate automated tracking performance of our tracking + cell division correction method on complete trajectories, individual manually annotated tracks from a complete dataset (Figure S1B) were used as ground truth and compared against automated tracking results. Each manual track was associated with one or more automated tracks, and transitions between automated track IDs within a single manual track were counted as linking errors. The duration of each manual track and the average duration of its associated automated segments were computed to assess fragmentation. Overall metrics included the average number of linking errors per manual track, average track duration, and the global average duration of automated segments. Additionally, the number of manual and automated cell divisions was quantified, enabling comparison of division detection accuracy.

### Shape feature computation from point clouds and membrane masks

After converting each segmentation label into a binary mask and extracting its surface using scikit-image’s ‘marching_cubes’ algorithm [67], we obtain a triangulated surface mesh. The vertices of this mesh are treated as a 3D point cloud. From this representation, we extract several shape features, including surface area, eccentricity, and a characteristic radius.

To estimate the surface area of an object, we compute the convex hull of its surface point cloud using scipy.spatial ConvexHull function [68]. The convex hull is the smallest convex polyhedron that encloses all points in the cloud. The total area of the triangular faces of this hull provides a robust approximation of the object’s surface area. To quantify the anisotropy of the shape, we compute the covariance matrix of the 3D point cloud. This 3 × 3 matrix encodes the spread of points along each axis. Performing eigenvalue decomposition gives the principal directions and the variances along them:

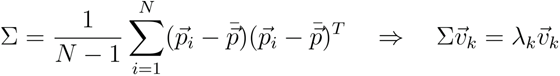

The square roots of the eigenvalues, referred to as eccentricities, measure the object’s spatial extent along each principal axis. Their relative magnitudes indicate how elongated or isotropic the shape is.

To obtain a scalar measure of object scale, we compute the geometric mean of the eigenvalues of the covariance matrix:

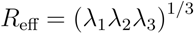

This value captures the volumetric spread of the object in all directions and is rotationally invariant. This effective radius provides a single, interpretable descriptor of the object’s overall size, and complements the more detailed eccentricity and surface area metrics.

Additionally, to get representative 2D measures from each membrane object, centroids were computed for each unique membrane object in the 3D dataset. The center slice – defined as the z-slice nearest to the rounded z-coordinate of the centroid – was cropped for each object. From these 2D center slices, morphological features including area, extent, eccentricity, and solidity were quantified using the ‘regionprops’ function from the scikit-image Python package[67].

### Nuclear tracking

To track segmented nuclei over time, we used the Fiji plugin TrackMate 7 [49], allowing us to input integer-labelled segmentation masks directly as objects using the Label Image Detector module. For temporal linking of spots, we applied the Linear Assignment Problem (LAP) tracker, with parameters optimized to accommodate our imaging frame rate and observed nuclear movement, specifically 16 µm linking max distance, 15 µm gap closing max distance and 3 max frame gap, disabling track splitting and merging, but applying TrackMate-Oneat branching correction (see details below).

The resulting track XML was used as input for our NapaTrackMater Python-based module, with which we calculate single-cell shape features for nucleus and membrane objects associated with the tracks, and calculate additional movement features based on nucleus tracking lineage information. For each channel, we calculated a “master XML” file, combining trajectory XYZT information with single-cell features. This file is used to store tracks and features, unwrapping them into result CSV files for plotting and further data analysis when needed.

For manual curation of TrackMate-generated tracks, we used Mastodon, an extension on the MaMuT [60] tracking platform based on the BigDataViewer plugin on Fiji [69]. We input the tracks on the dataset converted to an H5 file, and manually corrected cell lineages for as long as the cell was able to be visually followed. For backtracking, we input the manually annotated cell type XYZ locations (see details below) in the final frame of the Mastodon dataset, and assigned the manually curated trajectory of the corresponding final frame cell a cell type.

### Division detection with TrackMate-Oneat

We developed a pipeline leveraging deep learning-based action classification to detect cell division events and use these predictions to improve nucleus tracking. A key component of this system is the Oneat classifier, which was trained to distinguish mitotic from non-mitotic cells based on short temporal sequences of image data (Figure S7). For training, the 3D+t nucleus channel of a dataset was manually annotated for division events. Specifically, mitotic events are annotated on Napari by clicking on the center of the dividing nucleus in a microscopy time series. For each annotated division, a 64 x 64 x 8 voxel crop of 3 time frames was extracted around the clicked location, centered both spatially and temporally on the mitotic event. This creates a positive training sample. To create negative (non-mitotic) samples, a corresponding number of randomly selected locations are extracted from non-dividing nuclei. These negative and positive samples are used to train the model in a supervised fashion, optimizing a binary classification loss to distinguish mitosis from non-mitosis using a DenseNet-based architecture [70] (Figure S7).

For the prediction of division events from whole datasets, the trained Oneat model processes each object identified from a pre-generated nucleus segmentation. For each segmented nucleus at every time point, a temporal window is constructed by cropping 64 x 64 x 8 voxels for 3 frames centered on the nucleus XYZ centroid. These volumes are passed through the Oneat model, which classifies each central frame as either mitotic or non-mitotic.

To combine division events with tracking data of a dataset, the TZYX coordinates of predicted mitotic nuclei were recorded in a CSV file, which was used as input for the TrackMate-Oneat extension of TrackMate. This step uses the predicted division locations to impose a branching point on a trajectory. Suitable daughter cells for mitotic cells are searched from a 16.5 µm search radius from the mother cell, and linking is optimized using a Jaqaman linker algorithm [71]. This biologically-informed relinking improves the completeness and accuracy of lineage tracking, especially in datasets with frequent cell divisions.

To avoid spurious links and ensure geometric plausibility, the pipeline also incorporates the Mitosis Angular Region of Interest (MARI) principle. This constraint limits the search for daughter cells to a angular region from the mother cell’s nucleus principal axis of a fit ellipsoid. Candidate daughter spots were defined as those within a radial distance *r* of the mother spot m = (*x*_*m*_, *y*_*m*_) whose displacement vector s - m, with s = (*x*_*s*_, *y*_*s*_) the candidate spot position, formed an unsigned angle

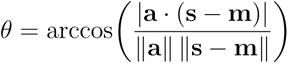

with the mother’s principal axis a not exceeding a threshold *λ*_max_ set by the user.

By restricting candidate daughters to fall within a defined angular region of interest, this method eliminates improbable pairings and enhances the biological realism of the reconstructed lineages. This constraint is especially important in dense tissues, where purely distance-based linking may result in incorrect associations.

### Cell final fate labelling

To annotate the cell types at the end based on actual cell fates, the final frame of live acquisition and the 3D acquired image of the IF-stained sample were aligned using the BigWarp tool in the BigDataViewer ImageJ plugin. Registration points were found by comparing nuclei channels (H2B for live image and DAPI for IF image). The warp settings were then applied to all channels of the IF image. Cell type XYZ locations were then manually annotated using the Napari Points tool using the logic in Table 1. (Fig. 5A)

#### Principal Components Analysis (PCA)

To reduce dimensionality and visualize feature trends, we performed Principal Components Analysis (PCA) using the ‘sklearn.decomposition.PCA’ implementation from the scikit-learn Python package. For each feature category (membrane shape, nucleus shape, position, movement, and all features), standardized feature values were extracted. PCA was fit and transformed on the selected features, and the resulting principal components (PC1 and PC2) were used for downstream analyses and visualization.

Feature contribution to the principal components of each PCA fitting was assessed by calculating the Spearman rank correlation between each feature and the two principal components (PC1 and PC2) from the corresponding feature category PCA. The correlation matrix was computed using ‘pandas.DataFrame.corr(method=’spearman’)’[72], and features were ranked by the absolute value of their correlation with PC1. The top-ranked features per category were selected as representative for further visualization and interpretation.

### Feature-time correlation analysis

For each cell trajectory, standardized features were correlated with experimental time. Tracks were included if their time span was at least 1 h to ensure sufficient temporal variation. For each track, Spearman’s rank correlation coefficient (ρ) was calculated between each standardized feature and the corresponding time in hours, using ‘scipy.stats.spearmanr’ from SciPy Python package.

The resulting per-track correlation coefficients for each feature were aggregated across tracks to obtain a distribution of ρ values per feature. To assess whether features were, on average, significantly correlated with time, a Wilcoxon signed-rank test was performed on raw ρ values. P-values were adjusted for multiple testing using the Benjamini–Hochberg false discovery rate (FDR) procedure. Correlation distributions were visualized as boxplots and statistical significance was indicated by asterisks placed above the corresponding feature’s box, denoting features with FDR-corrected p < 0.001.

### Feature separability scores

To quantify the separability of cell types based on PCA embeddings, several metrics were computed for each time point (Table 2).

**Table 2.** Separability metrics for feature category comparison.

| Metric name | Definition |
| --- | --- |
| KNN accuracy | The classification total accuracy of a k-nearest neighbors classifier (`'sklearn.neighbors.KNeighborsClassifier'`) trained to predict cell type from PC1 and PC2. |
| KMeans ARI | The agreement between cell type labels and k-means clustering assignments (``sklearn.cluster.KMeans``), measured using ``sklearn.metrics.adjusted_rand_score``. |
| KMeans NMI | The mutual information between cell type labels and k-means clusters, using ``sklearn.metrics.normalized_mutual_info_score``. |
| ASW | The silhouette score for cell type clusters in PCA space, using ``sklearn.metrics.silhouette_score``. |

These metrics were calculated for each feature category and visualized over time to assess how well cell types could be separated in the reduced PCA space.

### Data pre-processing for classification

Prior to classification, cell trajectory data were filtered to retain only timepoints with complete feature and cell fate information. A curated set of non-redundant features (Pearson cross-correlation < 0.9) was selected from the measured features. Features were standardized using z-score normalization. Rows with missing values were dropped. For model training and testing, a single dataset was used for the 5-class XGBoost or logistic regression classifier (see details below). In both cases, tracks were split such that no cell (and its corresponding track across time) appeared in both training and testing sets.

For the XGBoost model, class imbalance was addressed using a combination of undersampling (basal and goblet classes) and SMOTE-based oversampling of ionocyte and SSC classes. Undersampling was stratified over time by dividing the continuous time variable (‘t_hours’) into 10 quantile-based bins (each around 2 hours), ensuring that the temporal distribution of samples was preserved as much as possible. Within each bin, class samples were drawn proportionally so that each class contributed an equal number of examples overall.

Subsequently, the minority classes were oversampled using ‘SMOTENC’ class from the imbalanced-learn Python package [73,74]. SMOTENC (Synthetic Minority Over-sampling Technique for Nominal and Continuous features) extends the SMOTE algorithm to handle datasets with both categorical and continuous variables. It generates synthetic samples for minority classes by linearly interpolating between a true sample and its nearest true neighbors in feature space for continuous variables, using the binned time as a categorical variable. Oversampling was performed until the number of synthetic samples in each minority class matched the undersampled target size, yielding a fully balanced dataset across all five cell types.

### Single cell fate prediction

To predict cell fate from individual timepoints, we trained a multiclass XGBoost classifier using single-cell features extracted from each frame of every tracked cell. Each frame was treated as an independent sample and labeled by the final immunostaining cell type of its parent track. Preprocessed feature vectors were used as the independent variables for training, with experimental time (‘t_hours’) included as an additional feature for prediction.

For model training, the dataset was split into training and testing sets in a stratified manner across both timepoints and cell types. Class imbalance was addressed as described above. Model performance was evaluated using accuracy (*ACC*) and balanced accuracy (BACC), where

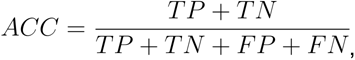

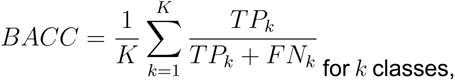

and a confusion matrix showing per-class true/false positive percentages. All comparative calculations were performed by comparing predicted and ground truth single observations’ cell type labels, using ‘accuracy_score’, ‘balanced_accuracy_score’, and ‘confusion_matrix’ classes from from scikit-learn Python package, and averaged across 20 independent iterations with random resampling to ensure robustness. Additionally, prediction confidence was estimated using the class probabilities output by the XGBoost model via the ‘predict_proba’ function. For each cell, we recorded the predicted class probability (i.e., the model’s estimated probability of its top prediction). These confidence scores were then averaged by cell type and time point within each iteration, followed by averaging across all iterations. Finally, a 1-hour rolling average was applied to visualize temporal trends in prediction confidence.

To assess the contribution of individual features to the XGBoost model, we performed an analysis leaving a single feature or whole feature categories out by iteratively retraining the model with each feature / feature category removed and measuring the change in accuracy metrics. This was also applied iteratively across 20 independent iterations, taking the mean change in balanced accuracy as the metric to sort the features.

For comparison, we also trained a logistic regression classifier using the same input data, using the ‘LogisticRegression’ class from the scikit-learn Python package. This model was implemented with L1 regularization and trained using the ‘saga’ solver with a maximum of 1000 iterations. The logistic regression results were used for comparison with a classwise confusion matrix, additionally, feature weights were output from the trained model to assess feature importance. Model performance assessment was performed across 20 iterations and averaged across iterations, similarly to XGBoost performance assessment.

## Acknowledgements

We thank reNEW Imaging Platform members for technical support and help with imaging, L. Davidson (University of Pittsburgh) for the mem-mNeonGreen pCS2 plasmid. We also thank members of the Sedzinski and Dumitrascu groups for comments and suggestions. This work was supported by grants from Novo Nordisk Fonden (NNF21CC0073729, J.S.), European Research Council Consolidator Grant (ERC CoG 101125803 MechanoFate, J.S.). M.T. acknowledges the support of the Copenhagen Bioscience PhD Program (NNF19SA0035442). J.S. acknowledges the support of the Novo Nordisk Foundation (NNF22OC0076414, NNF19OC0056962) and Leo Foundation (LF-OC-19-000219). B.D. acknowledges support by NIH Grant 1R35GM157082-01; B.D. holds a CIFAR Macmillan MHU Fellowship. This work was performed using HPC resources from GENCI-IDRIS (A0161013396, AD011013695).

## Disclosure and competing interests statement

V.K. is the founder of a not-for-profit company, Kapoorlabs, which was compensated for analysis work related to this study. The company operates on a not-for-profit basis and has no commercial interest in the outcomes of this research. All other authors declare no competing interests.

## Declaration of generative AI and AI-assisted technologies

While preparing this work, the authors utilized ChatGPT to assist with rephrasing certain sentences. All content generated with the tool was subsequently reviewed and edited by the authors, who take full responsibility for the final version of the publication.

## Data availability

### Lead contact

Requests for further information and resources should be directed to and will be fulfilled by the lead contact, Jakub Sedzinski.

### Materials availability

This study did not generate new materials.

### Data and code availability

Source data statement: Microscopy data reported in this paper will be shared by the lead contact upon request.

Code statement: Code for image data processing workflow is found at https://github.com/kapoorlab/CopenhagenWorkflow.

Code for data postprocessing, plotting, and analysis is found at https://github.com/Sedzinski-Lab/trackomics/.

## Author contributions

Conceptualization: M.T., Z.X., O.B., V.K., B.D., J.S.; Methodology: M.T., Z.X., O.B., V.K.; Software: M.T., Z.X., O.B., V.K. Validation: M.T, V.K.; Formal analysis: M.T, Z.X, O.B, V.K; Investigation: M.T.; Resources: M.T., V.K. J.S.; Data curation: M.T., V.K.; Writing - original draft: M.T., J.S.; Writing - review & editing: M.T., Z.X., O.B., V.K., B.D., J.S.; Visualization: M.T., Z.X., O.B., V.K.; Supervision: B.D., J.S.; Project administration: M.T, J.S; Funding acquisition: J.S.

---

**Figure S1.**
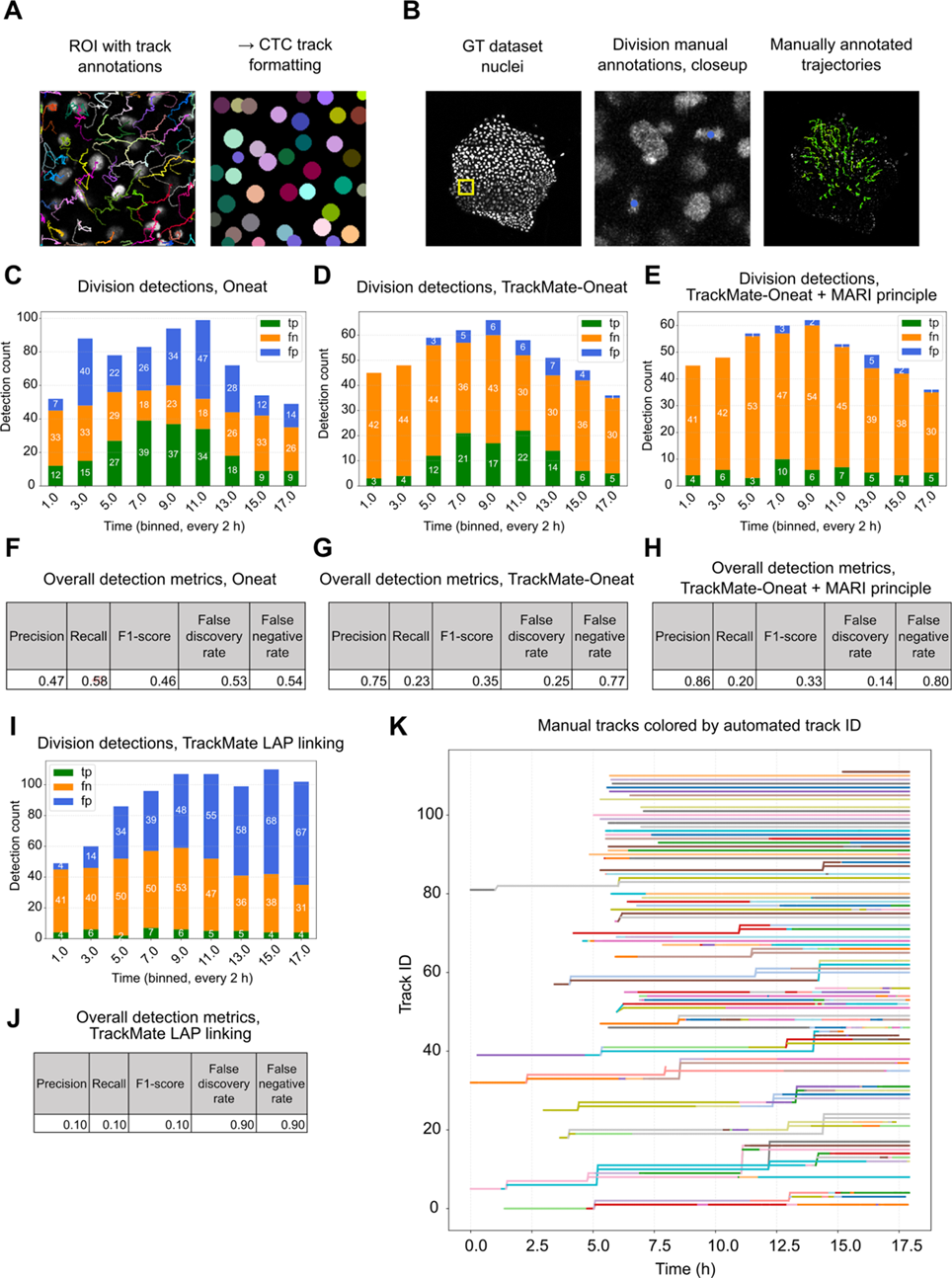
Quality metrics for tracking and division detection. **A-B.** Cell Tracking Challenge format quality estimations in Table S3 are based on an ROI of a dataset (A), where each cell is manually tracked for as long it appears in the ROI. Manual track annotations are formatted in CTC format, to which automated tracking with TrackMate is then compared to estimate tracking quality. Division metrics in C-J are based on a dataset where each cell division is manually annotated (B). In this dataset, selected tracks are also annotated and compared to automated tracking in Table S3 and K. **C-E, I.** Division detections for Oneat (not connected with tracks), TrackMate-Oneat (Oneat divisions connected with TrackMate tracks), TrackMate-Oneat + MARI principle (TrackMate-Oneat with max boundary set for angle between mother cell and daughter cells), and TrackMate “native” track splitting, enabled in TrackMate LAP linking algorithm. **F-H, J.** Corresponding detection metrics. **K.** Manually annotated ground truth tracks colorized by the Track ID assigned by automatic tracking used for the experiments.

**Figure S2.**
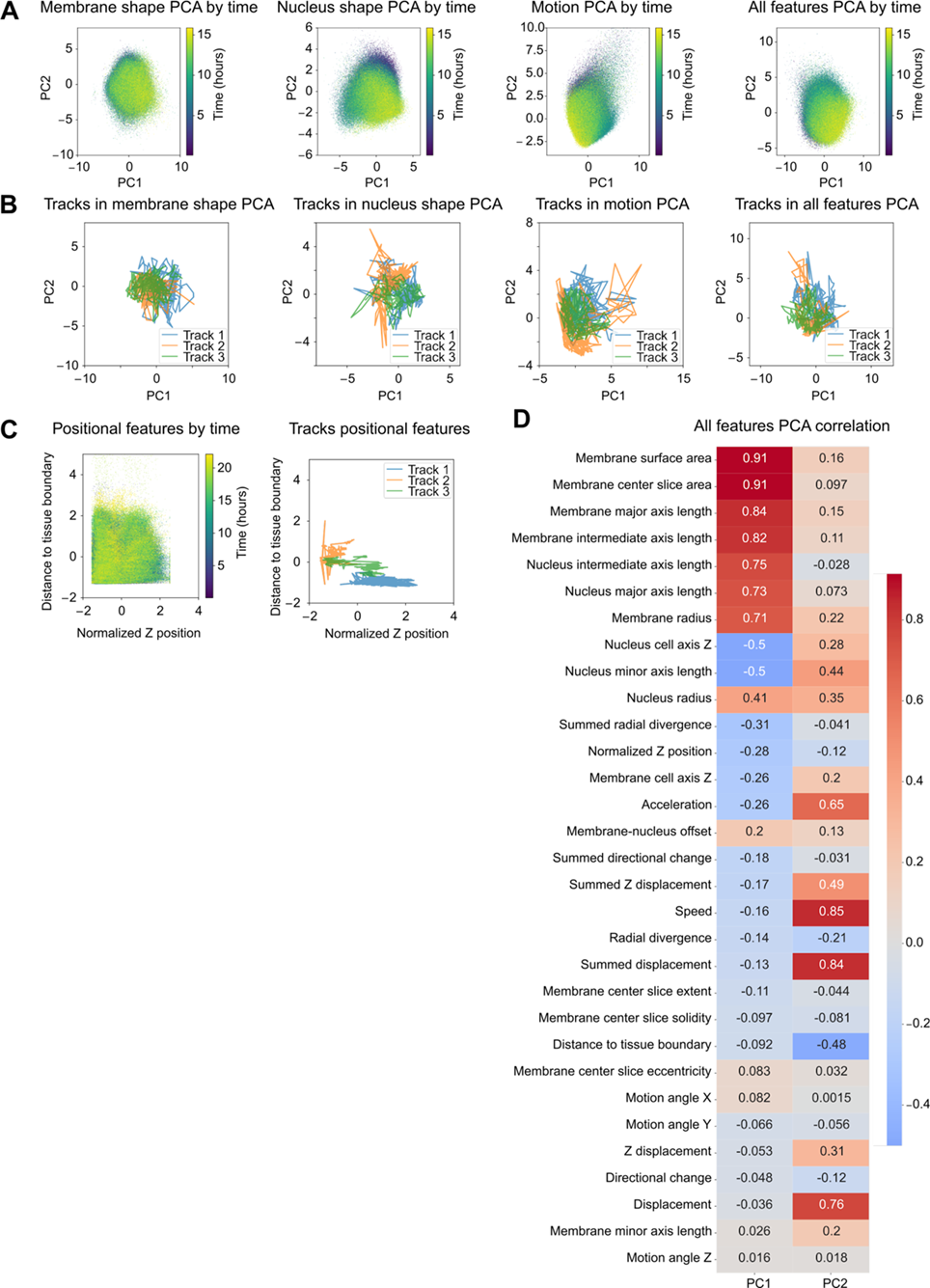
Supplemental data for principal component analysis. **A.** PCA feature spaces of dataset 1. **B.** Individual cell trajectories in feature spaces fit for dataset 2. **C.** Timewise positional features and individual trajectories’ positional features for dataset 2. **D.** All features’ correlation with all feature PC1 and PC2.

**Figure S3.**
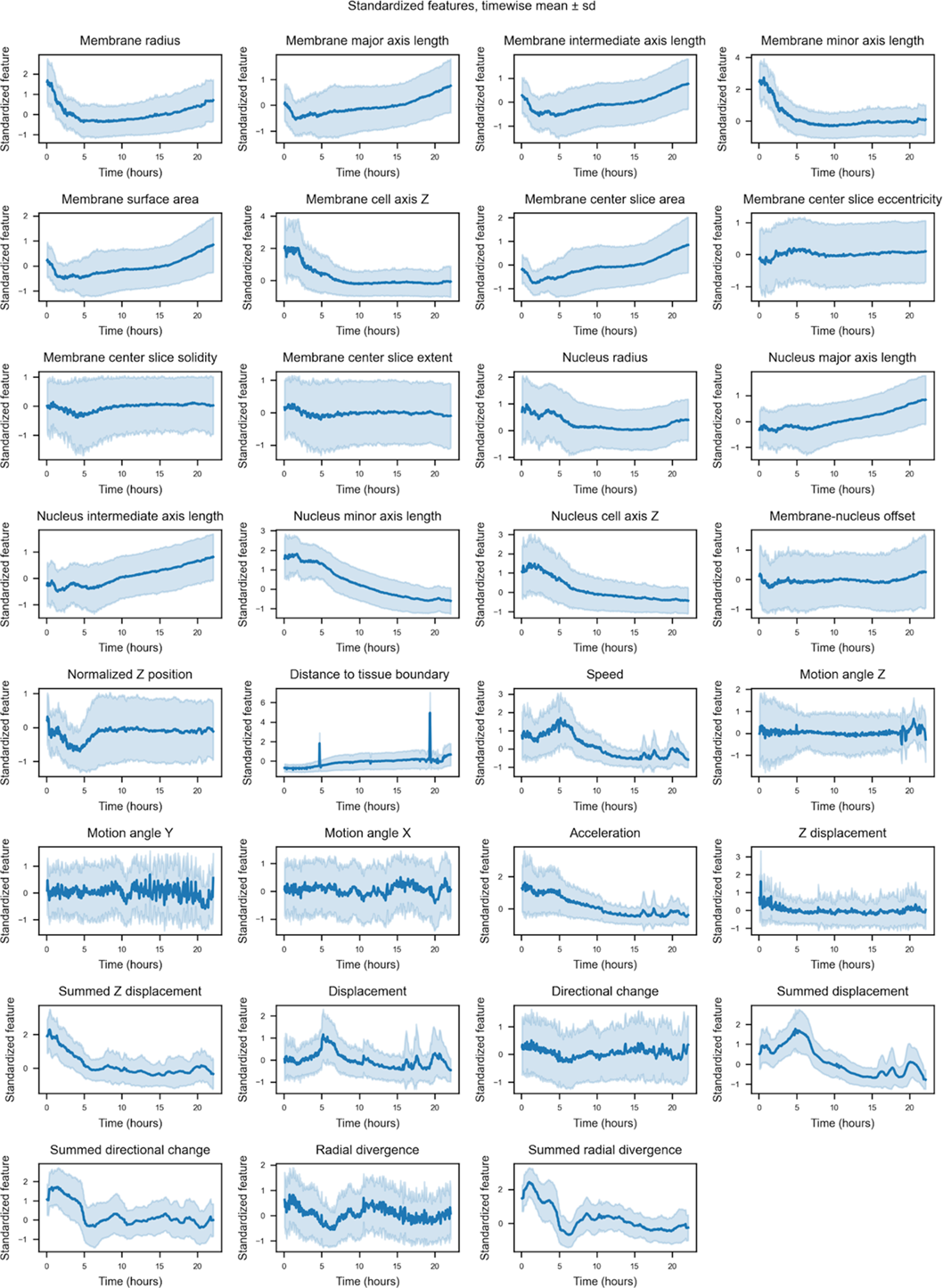
Mean and s.d. of all features across time, dataset 2.

**Figure S4.**
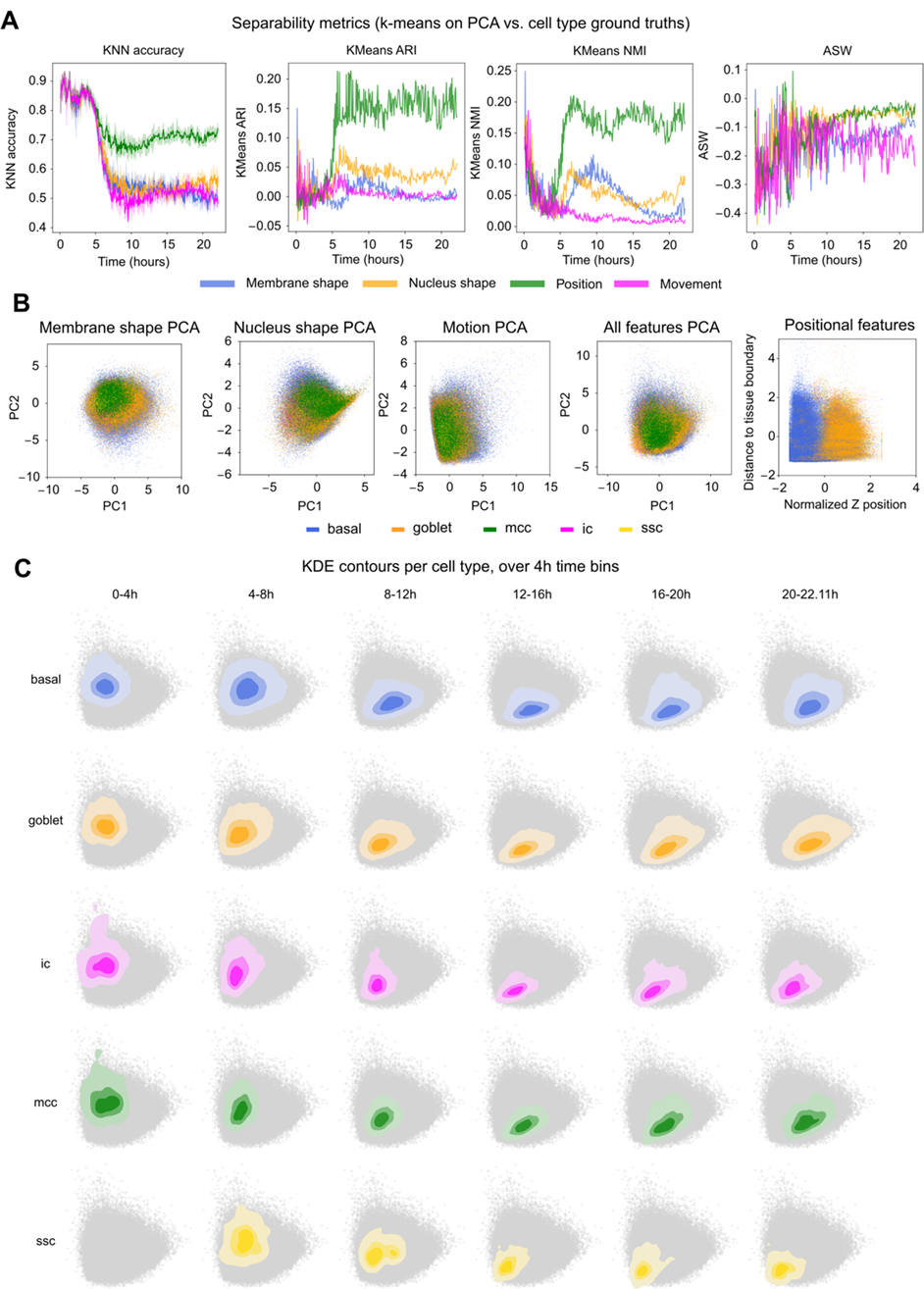
Backtracked cell types in PCA feature space. **A.** Separability metrics for feature categories. Mean accuracy score (KNN accuracy) based on KNearestClassifier (scikit-learn) of different feature categories where k = 5, Adjusted Rand Index score (KMeans ARI), Normalized Mutual Information score (KMeans NMI) for comparing k-means clustering of selected feature spaces (corresponding to panel B) with ground truth cell type labels. Average silhouette width (ASW) is calculated for different feature categories against ground truth cell type labels. **B.** Cell types labelled in feature spaces corresponding to Fig. 3D and Fig. S2C. **C.** Kernel density estimation of time windowed cells in the all features PCA space, estimated per cell type.

**Figure S5.**
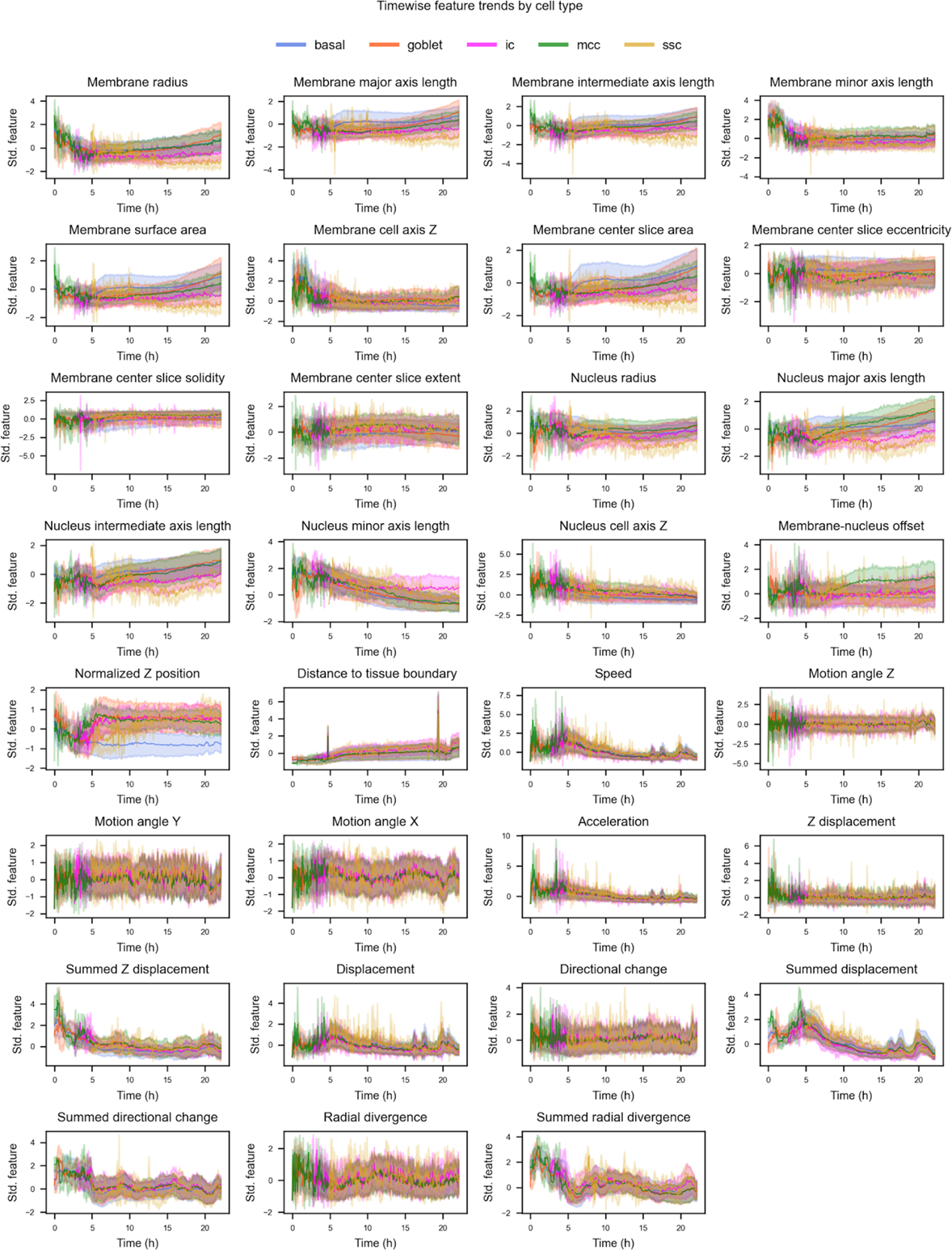
Mean and s.d. of all features of cell type annotated cells across time, colored by cell type, dataset 2.

**Figure S6.**
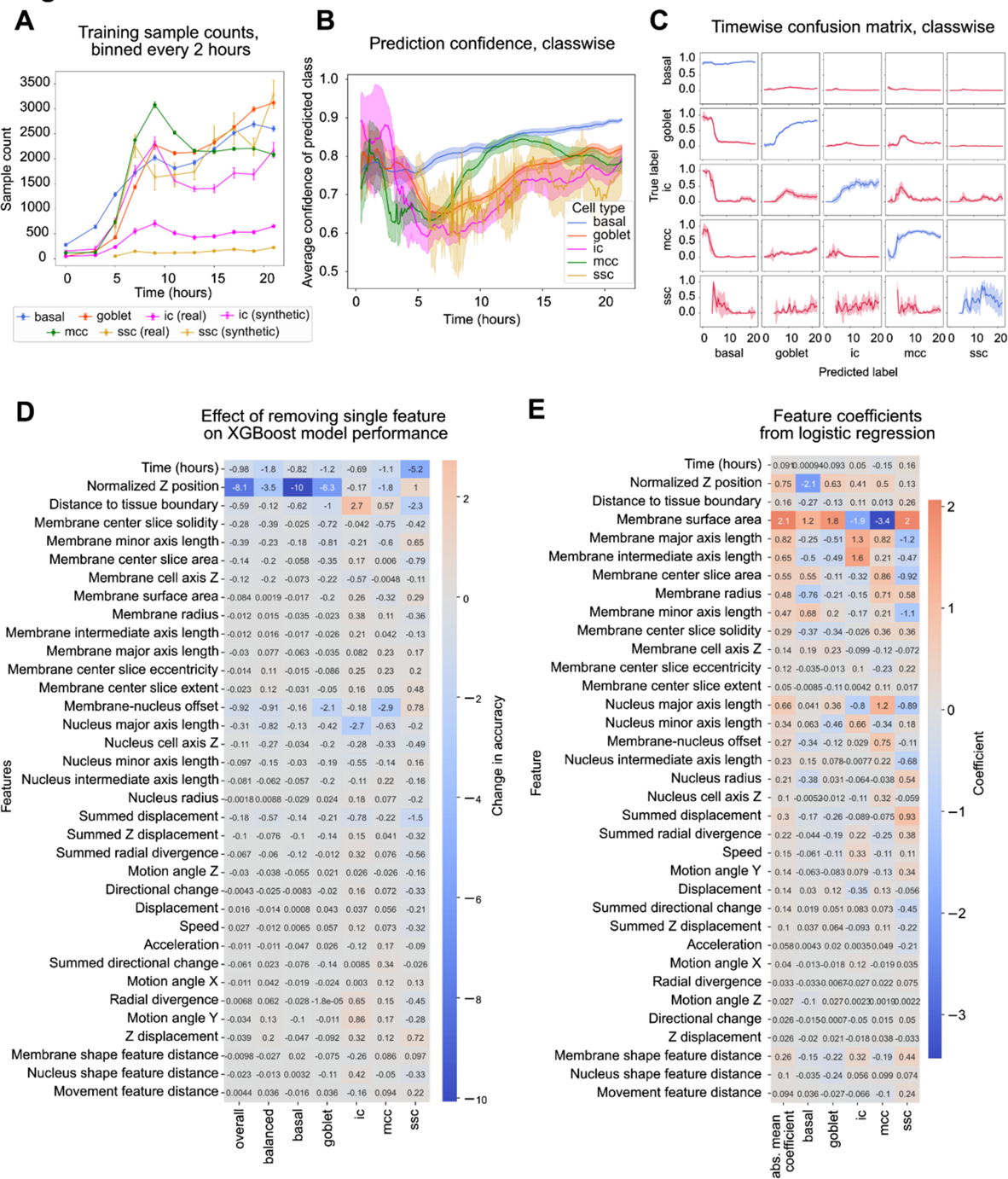
Classifier model performance analysis and feature importances. **A.** Classifier training dataset sample counts across dataset time. For basal, goblet, and mcc classes, only real samples were used, for ic and ssc classes, some samples were synthesized using SMOTE oversampling (imbalanced-learn). **B.** Mean and s.d. prediction confidence of predicted class of XGBoost-based predictions, colored by class. **C.** Timewise accuracy confusion matrix of XGBoost-based predictions. **D.** Percent point-wise accuracy difference to baseline accuracy of the XGBoost model for leaving out a single feature. **G.** Feature coefficients for a logistic regressor model. All calculations in B, C, D, and E are averaged out for 20 iterations of randomized test-train split, model training and prediction.

**Figure S7.**
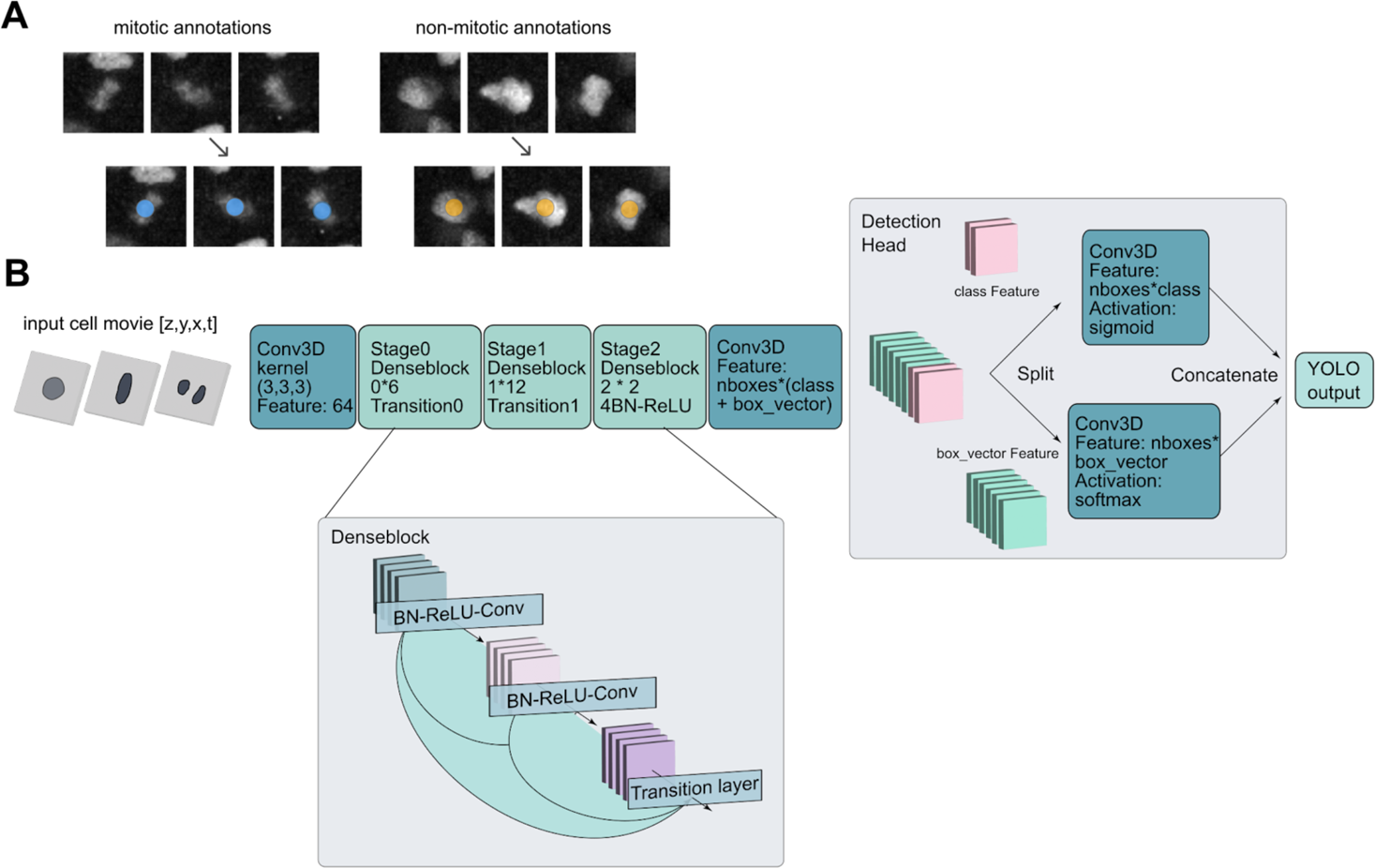
Oneat model structure. **A.** Training data annotations in Napari for training a mitosis classifier. **B.** CNN architecture. Input data is 3 timepoints, 8 x 64 x 64 pixel crop, centered around the annotation ZYX+t centroid. Output is probabilities for classification as mitotic or non-mitotic.

## Supplementary videos

**Movie S1. Timelapse movie of nucleus and membrane signals, 0-22 h.** Maximum intensity Z projection of raw nucleus (H2B-RFP) and denoised membrane (mem-mNeonGreen) signals from Dataset 1. Scale bar: 500 µm.

**Movie S2. Timelapse movie of 3D StarDist nucleus segmentation and 3D CellPose segmentation, 0-20 h.** Maximum intensity Z projection of raw nucleus and denoised membrane signals together with integer labelled segmented nuclei and membrane segmentation results from StarDist and CellPose from Dataset 3. Scale bar: 500 µm.

**Movie S3. Combined membrane segmentation, nucleus segmentation, and tracking data for a single cell, 0-16 h.** Left: 3D Napari rendering of the membrane and the nucleus segmentations isolated for a single tracked cell, overlaid on a cropping their respective channels. Right: corresponding track from TrackMate, visualised in Napari, overlaid on maximum Z projection of the denoised membrane channel, from Dataset 2.

**Movie S4. Oneat division detection pipeline.** Annotation: dividing cells and non-dividing cells are manually annotated in Napari by observing the nucleus channel timelapse and making Point annotations on the approximated center of the cell. Training data: The model is trained from TZYX crops of (3, 8, 64, 64) centered around the annotation coordinate. 3 training patches for dividing cell and normal cell are depicted here as a maximum intensity Z projection. Division prediction: Oneat produces predictions of division TZYX coordinates, visualised in Napari by selecting StarDist segmented objects that overlap with the division coordinates. Scale bar: 500 µm.

**Movie S5. Timelapse movie of tracking results with nuclei, 0-20 h.** Maximum intensity Z projection of raw nucleus signal together with trajectories from TrackMate, visualised using Mastodon ImageJ plugin, from Dataset 3. Trajectories shorter than 5 timepoints were left out from the visualisation. Scale bar: 500 µm.

**Movie S6. Backtracked cell types’ trajectories, 0-22 h, colored by cell type.** Visualisation of trajectories with a cell type label based on end timepoint annotation, overlaid on maximum intensity Z projection of raw nucleus signal. MCC, IC and SSC trajectories are all manually curated, most basal and goblet trajectories are automatically tracked and assigned a cell type based on their end timepoint cell type annotation. From Dataset 1. Scale bar: 500 µm.

**Movie S7. Selected individual cells’ cell shapes across 0-22 h, colored by cell type.** 3D Napari rendering of membrane objects of 5 selected single-cell trajectories per cell type. If no segmented object is found for a timepoint, the previous valid segmentation is shown.

## Notes

### Summary of Updates

Added missing ORCID numbers and funding agencies

